# Gut microbiome changes over the course of multiple sclerosis differentially influence autoimmune neuroinflammation

**DOI:** 10.64898/2026.02.23.707375

**Authors:** Amélia Sarmento, Tommy Regen, Daniela Ferro, Ángel Ruiz-Moreno, Cintya González-Torres, Katlynn Carter, Arthi Shanmugavadivu, Ari Waisman, Maria José Sá, Carles Ubeda

**Affiliations:** FP-I3ID, Faculdade de Ciências da Saúde, Universidade Fernando Pessoa, 4200-150 Porto, Portugal; RISE-Health, Universidade Fernando Pessoa, 4200-150 Porto, Portugal; I3S, Instituto de Investigação e Inovação em Saúde, Universidade do Porto, Porto, Portugal; Institute for Molecular Medicine, University Medical Center of the Johannes Gutenberg University Mainz, Mainz, Germany; Research Center for Immunotherapy (FZI), University Medical Center of the Johannes Gutenberg University Mainz, Mainz, Germany; Serviço de Neurologia, Hospital Universitário de São João, Porto, Portugal; Departamento de Neurociências Clínicas e Saúde Mental, Faculdade de Medicina, Universidade do Porto, Porto, Portugal; Fundación para el Fomento de la Investigación Sanitaria y Biomédica de la Comunitat Valenciana - FISABIO, Valencia, Spain; Centers of Biomedical Research Network (CIBER) in Epidemiology and Public Health, Madrid, Spain

## Abstract

Multiple sclerosis (MS) is the leading inflammatory and demyelinating disease of the central nervous system (CNS). MS begins with systemic inflammation and over time is compartmentalized within the CNS. Studies in humans, supported by animal experiments, suggest that the gut microbiome plays an important role in MS development. However, despite the dynamic nature of the disease, little is known on how the microbiome evolves over the course of MS and how these microbiome changes influence immune responses and disease progression. Here, using high-throughput sequencing, we identified distinct gut microbial communities with differential functional potential in MS patients stratified by time since disease onset. Importantly, using a humanized mouse model, we demonstrate that differences in microbial composition significantly impact disease outcomes. Microbiota from more recently diagnosed MS patients induced severe neuroimmune disease in mice, whereas microbiota from long-term MS patients and healthy individuals elicited only mild disease. Accordingly, we found that microbiota from earlier diagnosed MS patients exhibit a higher inflammatory potential. This was characterized by a reduced capacity to induce regulatory T cells in mice and an increased induction of pro-inflammatory cytokines in human peripheral blood mononuclear cells. Together, our results suggest that the ability of the gut microbiome to promote systemic inflammation and trigger MS pathology shifts over the course of the disease and is primarily critical during its early stages. These findings indicate that a limited therapeutic window should be considered when designing microbiome-based interventions for MS.

## INTRODUCTION

Multiple Sclerosis (MS) is a chronic autoimmune demyelinating disease of the central nervous system of unknown aetiology, that develops when the immune system erroneously recognizes and attacks the myelin covering of the nerve fibres, leading to focal inflammation and axonal/neuronal damage. Clinically, two major phenotypes are nowadays recognized, relapsing or progressive, each of them further subdivided according to disease activity (Lublin et al 2014). The vast majority of MS patients develop a relapsing form, with episodes of acute disease activity followed by periods of complete or partial remission in the first years of the disease, corresponding to the classical relapsing/remitting MS subtype – RRMS. With time, a considerable number of RRMS patients experience a progressive accumulation of disability, referred to the classical secondary progressive subtype (SPMS) (Pontieri et al., 2024; Zanghì et al., 2025).

Importantly, MS is a dynamic disease in which the mechanisms driving the inflammatory and neurodegenerative process change over time. Based on pathophysiological mechanisms, MS progression has been postulated to occur in three stages (Leray et al., 2010; Lassmann, 2017). In a first stage, occurring primarily in the early phase of RRMS, brain injury is mainly mediated by systemic inflammation. This stage is characterized by systemic high levels of pro-inflammatory cytokines and recruitment of leukocytes from distal sites to the brain. In a second stage, occurring at late RRMS and early progressive MS, inflammation is more compartmentalized, and T and B cells are mainly localized within the brain. Finally, in the third stage, occurring at the late phase of progressive MS, inflammation is reduced and progressive neurodegeneration occurs even in the absence of an inflammatory response.

Although certain genetic polymorphisms predispose to MS (Baranzini and Oksenberg, 2017; Goris et al., 2022), studies performed with monozygotic twin pairs have suggested that environmental factors play a major role on disease initiation and progression (Fagnani et al., 2014). Within these factors, due to the impact of commensal gut microorganisms on immune responses, extensive attention has been paid to the potential role of the gut microbiome on MS progression. MS patients have been shown to carry distinct gut bacterial communities as compared to healthy individuals (Miyake et al., 2015; Jangi et al., 2016; Tremlett et al., 2016; Berer et al., 2017; Reynders et al., 2020; Horton et al., 2021). Moreover, the usage of mouse models has strongly suggested that some of the detected microbial changes do play a role in MS development. Indeed, germ-free (GM) mice inoculated with faecal microbiome from MS subjects were shown to contain lower number of Treg cells than those inoculated with the microbiome from healthy individuals, and subsequently developed a more severe disease (Cekanaviciute et al., 2017).

As described above, MS is not a static disease, therefore, it is plausible that the gut microbiome also evolves concomitantly with disease progression and that these microbial changes could impact immune responses and influence disease development. This hypothesis is partially supported by a few studies in which the abundance of different bacterial taxa has been shown to differ between different MS subtypes, i.e. RRMS vs SPMS (Cox et al., 2021; Zhou et al., 2022). However, it is not well understood whether these microbiome changes actively impact MS pathology, particularly with respect to the changing immune responses observed during MS progression. Moreover, although differences in the microbiome have been detected between MS subtypes, there are no studies that have investigated such differences within a single subtype over the course of the disease. This is particularly relevant in the RRMS subtype, in which, as described above, two potential stages have been proposed based on the switch from predominantly systemic inflammation (early phase) to a more localized inflammation (late phase).

In this study, we evaluated the gut microbiome of patients with RRMS of varying disease durations using high-throughput sequencing. We show that disease duration is associated with changes in the microbiome in these patients. Moreover, using a mouse model and human peripheral blood mononuclear cells, we demonstrate that the gut microbiome of RRMS subjects with a shorter disease duration is more prone to induce systemic inflammatory responses and markedly intensified disease severity in the experimental autoimmune encephalomyelitis (EAE) mouse model, as compared to human microbiomes from RRMS patients with a longer disease duration. Altogether these results suggest that the MS gut microbiome evolves and plays a differential role on MS over time, a result that will have implications on the design and implementation of microbiome-based therapies towards the resolution of MS.

## RESULTS

### Microbiome structure is associated with RRMS disease duration

Twenty-seven adult MS patients were recruited to analyze their microbiome composition. As control, fecal samples from a similar number of individuals not developing MS (N=26) were included in the study. Patient’s characteristics are described in **Suppl. Table 1**. Most patients were developing RRMS (N=25, 92.6%), while 2 of them had SPMS and were therefore excluded from subsequent analyses. All RRMS patients, hereafter we will refer to them as MS patients, were treated with disease modifying therapies (DMT, **Suppl. Table 1**): 88% started platform drugs (interferon beta, glatiramer acetate), while the remaining started oral drugs (dimethyl fumarate, fingolimod). Eleven patients on platform drugs also initiated the DMT fingolimod and/or natalizumab. Concomitant diseases were detected in most patients (N=16, 64.0%), being the most frequent one depression (N=12, 48.0%). The mean and median disease duration was 16 and 17 years, respectively. The mean age for MS patients was 44 and did not differ from controls (mean age: 46 years, *t-* test, *p* = 0.584). In the control group there was a slightly less proportion of females (53.8%) as compared to the MS group (76%), albeit non-significant (Fisher test, *p* = 0.144). Fecal samples were analyzed through 16s rRNA sequencing (see methods) and sequences were processed using DADA2 for amplicon sequence variants (ASVs) and taxonomic assignment. After applying quality thresholds, an average of 139,860 ± 30,759 sequences were included per sample, which led to the identification of an average of 414.16 ± 116.55 ASVs per sample. No differences in the number of obtained sequences were detected between MS and control subjects (*p* = 0.831, *t*-test). The identified ASVs represented for most samples the maximum number of ASVs that could be detected based on rarefaction curves, and based on comparison with the Chao Index (the potential maximum number of ASVs), which was almost identical to the real number of detected ASVs (**Suppl. Fig. 1**). Thus, the obtained sequencing depth was sufficient to accurately determine the fecal microbiota composition in our patient’s cohort. We first analyzed overall differences in the microbiome composition that could be associated with disease status. No significant differences in alpha diversity − measured by both richness and Shannon diversity index − were detected between MS and non-MS individuals (**Fig. 1A**; *p* > 0.32, *t-*test). Similarly, no differences in beta diversity were detected between the two groups (**Fig. 1B**; PERMANOVA, *p* = 0.223). These findings are consistent with a previous report showing no significant differences in overall microbial diversity between MS and healthy individuals (Berer et al., 2017). However, a previous study, based on a larger cohort, reported significant differences in beta-diversity associated with MS (Zhou et al., 2022). Beyond the relatively small size of our cohort, another potential explanation for the absence of detectable differences in microbiome structure between MS patients and control subjects could be the heterogenous composition of both groups under study. In fact, we notice that 50% of the control individuals presented common non-neurological conditions, mainly hypertension (N=4, 15.4%), dyslipidemia (N=4, 15.4%) or rhinitis (N=3, 11.54%) (**Suppl. Table 1**). We first evaluated if these conditions had any effect on the microbiome composition that could affect subsequent comparisons with MS patients. Based on a Principal Coordinate of analysis (PCoA) and PERMANOVA analysis, the microbiome of control individuals did not differ according to the presence/absence of these medical conditions (**Suppl. Fig. 2A**; PERMANOVA, *p* = 0.428). Consistently, beta diversity between MS patients and controls remained non-significantly different after excluding from the analysis control subjects with other medical conditions (**Suppl. Fig. 2B**; PERMANOVA, *p* = 0.709). Next, we examined if the heterogeneity within the MS population was associated with variability in microbiome composition, potentially limiting our ability to distinguish MS patients from controls. Patients receiving therapy have been found to carry a microbiome distinct from non-treated MS patients (Zhou et al., 2022). However, all patients in our cohort were treated, thus the absence of treatment was not considered a factor that could impact microbiome composition in our cohort of MS patients. Among other variables, neither age, nor gender, physical exercise, first relapse type, vitamin D deficiency, vitamin D supplementation, Expanded Disability Status Scale (EDSS) or concomitant depression were found to be associated with microbiome composition in our MS patient’s cohort (**Fig. 1C**; *p* > 0.05). Given the substantial variability in disease duration among our MS patients (min=2, max=42, SD=10.26 years) and the inherently dynamic pathophysiological nature of MS, we analyzed if disease duration could be associated with the microbiome composition. To this end, we divided the group of MS patients in two groups, according to the median disease duration (16 years). Remarkably, a significant difference in microbiome composition was found in MS patients according to disease length (**Fig. 1C**). Patients with a disease duration equal to or greater than the median (≥16 years), referred to as the long-MS group, exhibited a significantly distinct microbiome composition (**Suppl. Fig. 3A**; PERMANOVA, *p* = 0.01) than patients with a more recent disease onset (<16 years, short-MS). Thus, disease duration was the analyzed factor most significantly associated with differences in the microbiome composition in our cohort of MS patients.

**Figure 1.**
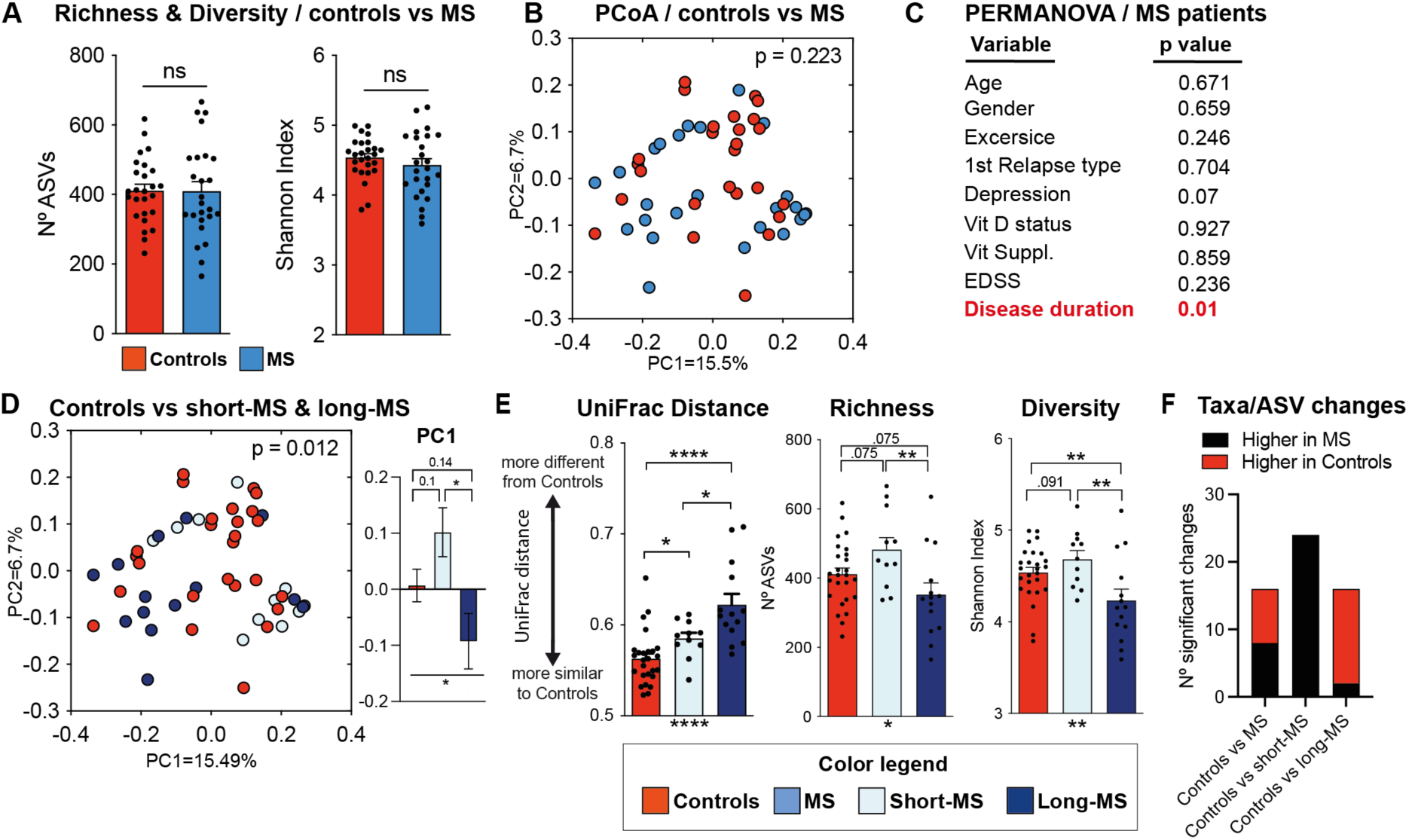
Changes in gut microbial communities are associated with MS duration. **(A)** Alpha-diversity (richness and Shannon index) does not significantly differ between MS and control subjects. **(B)** Beta-diversity, using the unweighted UniFrac distance, does not significantly differ between control and MS subjects. **(C)** The overall microbiome composition is significantly associated with disease duration (PERMANOVA test using unweighted UniFrac distances). **(D)** Beta-diversity of MS subjects significantly differ from controls (PCoA and PERMANOVA test based on UniFrac distances) when MS patients are stratified according to disease duration into short-MS (<16 years from onset) and long-MS (≥16 years from onset). These 3 groups are differentially distributed along the first principal coordinate (PC1) axes of the PCoA. **(E)** UniFrac distance from controls is significantly higher in long-MS than in short-MS patients. In addition, richness and diversity significantly differ among short-MS, long-MS and control individuals. **(F)** Most of the taxonomic changes between control and short-MS subjects consist on higher abundance of taxa in short-MS patients, while taxa depletion in long-MS patients is the most common change detected when compared with controls. Specific taxonomic changes (Boruta & ANCOM-BC, p<0.05) are shown in Suppl. Fig. 4. *p/q<0.05, **p/q<0.01, ****p/q<0.0001; T-test in (A); PERMANOVA test in (B-D); Kruskal-Wallis (below) and FDR for multiple comparisons (above) in (E, UniFrac) and One-Way ANOVA (below) and FDR for multiple comparisons (above) in (E, Richness and Diversity). When FDR test is applied, q values instead of p values are indicated. The exact q values are shown in (D and E) when q is between 0,05 and 0,1. Bars in A, D and E represent the average while whiskers represent SEM. N=11-26 human subjects.

### RRMS duration is associated with changes in microbial diversity and richness

Taking into account that disease duration was significantly associated with microbiome composition, we decided to stratify MS patients according to time elapsed since disease onset and reanalyzed their differences in microbiome composition compared to control subjects. In contrast with the previous results (**Fig. 1B**), the microbiome composition from controls was significantly different from MS patients when these were stratified according to disease length (**Fig. 1D**; PERMANOVA, *p* = 0.012). Accordingly, the first principal coordinate of analysis, explaining 15.49 % of the variability among samples, significantly segregated samples according to their MS status and time from onset (Kruskal−Wallis test, *p* = 0.017). To further understand how disease length could impact microbiome composition, we calculated and plotted UniFrac distances (measure of microbiome dissimilarity) between control subjects and short-MS and long-MS patients. Significantly higher UniFrac distances were observed between MS patients and control individuals compared to the distances among control subjects themselves (**Fig. 1E**, Kruskal–Wallis test & False discovery rate - FDR, *q* < 0.05). This result suggests that the microbiome of both MS subgroups (short- and long-MS) differs from that of controls, although the differences between long-MS patients and controls were significantly greater than those between short-MS patients and controls. Having shown that beta diversity differs between controls and MS individuals when these were stratified by disease length, we analyzed if the change in microbiome alpha diversity was also associated with MS disease duration. Remarkably, both Shannon diversity and richness significantly differ between short-MS and long-MS patients (**Fig.1E**; ANOVA & FDR, *q* < 0.01). Accordingly, an opposite change in alpha diversity was detected between controls and MS patients of different disease duration. A reduction in richness and Shannon diversity was detected in long-MS patients as compared to controls (**Fig. 1E**; ANOVA and FDR, *q* = 0.075 and 0.007 respectively), while short-MS patients had a higher microbiome alpha diversity (both in richness and Shannon diversity) than controls (**Fig. 1E**; ANOVA and FDR, q=0.075 and 0.091 respectively). Accordingly, the richness and Shannon diversity from short-MS patients was significantly higher than that one from long-MS individuals (**Fig. 1E**; ANOVA and FDR, q<0.01). In summary, within our patient’s cohort, the microbiome of MS patients differs from that of controls. However, these differences are only evident when patients are stratified by disease duration — a variable that is significantly associated with changes in microbiome composition and diversity during MS.

### RRMS duration is associated with depletion in specific taxa

Next, we analyzed, at every taxonomic level, microbiome differences associated with MS. These analyses were performed considering the whole cohort of MS patients and stratifying MS subjects according to disease duration. To this end, we applied a machine learning approach (Boruta algorithm) in order to identify those commensal bacteria with more power on differentiating MS individuals from controls, coupled with a univariate test (ANCOM-BC) to confirm the differences detected by machine learning. Following this approach, differences in 16 phylotypes, including 1 phylum, 1 class, 1 order, 3 families, 4 genera and 6 ASVs were detected between MS and control individuals (**Fig. 1F**, **Suppl. Fig. 4**; **Suppl. Table 2**, ANCOM-BC, *p* < 0.05). Increase of 7 phylotypes (43% of detected changes), mainly of the class Lentisphaeria, was detected in MS individuals, while 9 phylotypes, mainly from the class Clostridia, were depleted (56.25% of detected changes) as compared to controls. Next, we defined differences in MS individuals but stratifying them by disease duration. As expected, taxonomic changes detected in short-MS or long-MS individuals were remarkably different. Consistent with a lower richness and diversity in long-MS patients, most differences (14/16, 87,5%) detected in this group consisted on depletion of specific taxa, mainly from the class Clostridia and Methanobacteria (**Fig. 1F**, **Suppl. Fig. 4**; **Suppl. Table 2**, ANCOM-BC, *p* < 0.05). In contrast, and consistent with a higher richness and diversity detected in short-MS individuals, the 24 differences detected between short-MS and controls consisted on higher abundance of specific phylotypes on short-MS patients, mainly from the class Clostridia (**Fig. 1F**, **Suppl. Fig. 4**; **Suppl. Table 2,** ANCOM-BC, *p* < 0.05). Next, we identified taxonomic differences within MS individuals that were associated with disease length. The abundance of 34 phylotypes, including 2 phyla, 3 classes, 4 orders, 2 families, 5 genera and 17 ASVs significantly differ between short-MS and long-MS individuals (**Suppl. Fig. 3**; **Suppl. Table 2,** ANCOM-BC, *p* < 0.05). As expected, considering the higher richness and diversity detected in short-MS patients, the majority of changes (85%) consisted on higher levels of particular phylotypes in short-MS individuals. Thus, different taxonomical changes were detected in MS patients according to the length of the disease.

To validate our findings, we made use of a larger cohort that included 437 RRMS patients and household controls (Zhou et al., 2022) from four different countries within two different continents. Since all our patients were treated, we only included in our validation analysis a subset of 285 treated RRMS patients and their respective household controls. Notably, in this larger cohort we were able to recapitulate particular findings that were also detected in our patient cohort. Although microbiota richness and diversity did not differ between long-RRMS individuals and their respective household controls, microbiota richness was found to be significantly higher in short-RRMS individuals as compared to their respective controls (**Suppl. Fig. 5A**). Moreover, by applying the Boruta algorithm, we were able to identify several genera and ASVs that differ according to disease length (**Supp. Fig. 5B**; ANCOM-BC, *p <* 0.05). In concordance with the results obtained in our cohort, the majority of the differences detected consisted on the expansion of particular phylotypes of the class Clostridia on short-RRMS individuals. Thus, increase in the microbiota richness and higher abundance of particular phylotypes was found to be a common microbiome signature in RRMS patients with a shorter disease duration, both in our as well as in the validation cohort.

### Higher disability and depression are partially linked to microbiome changes associated with RRMS duration

Having shown that disease duration is associated with microbiome composition changes, we next attempted to identify variables associated with disease duration that could explain the microbiome changes detected. To this end, we first analyzed if the levels/prevalence of a particular variable was significantly different among the two MS stratified groups (short-MS, long-MS). Among studied variables (**Suppl. Table 3**), we found that patients with prolonged MS had significantly higher EDSS score (p=0.0085, Mann Whitney test), more prevalence of depression (p=0.015, Fisher test) and were, as expected, older (*p* = 0.0003, *t*-Test). We first focused on age as a variable that could explain some of the detected changes between short and long-MS patients. Age was not significantly associated with any of the phylotypes that significantly differ between short- and long-MS patients (Spearman correlation, p > 0.05). In contrast, within the 34 phylotypes that differentiated short and long-MS subjects, 9 of them were associated with EDSS score (**Suppl. Fig 6**). In all cases the detected phylotypes were depleted in patients with higher EDSS score, and as expected, were also depleted in long-MS patients which have a significantly higher EDSS score. Besides associations with EDSS, other taxa were found to be associated with depression. Specifically, the abundance of 16 taxa whose abundance significantly differ between long- and short-MS patients, were associated with depression (**Suppl. Fig 7**). The majority of the detected associations (87,5%) consisted on the reduction in the abundance of specific taxa in patients that suffer depression and in patients with longer disease duration, in which, as previously shown, concomitant depression was significantly more common. In summary, substantial microbiome differences were detected between short- and long-MS patients, and the majority of them (64,7%) were found to be associated with EDSS and/or depression, two variables that significantly differ between the two groups of MS stratified patients.

### RRMS duration is associated with changes in bacterial encoded functions

Having shown that MS duration is associated with changes in the microbiota, both at the taxonomic and diversity level, we next addressed if MS progression would also lead to changes in the microbiome functional potential. To this end, we performed shotgun metagenomic sequencing on the DNA extracted from previously analyzed samples. An average of 7.93×10^6^ ± 1.67×10^6^ high quality sequences were obtained per sample. No differences in sequencing coverage were detected between groups of samples (*t*-test, p=0.426). Bacterial encoded ORFs were identified and a total of 6,487 functional gene orthologs (5,304 ± 231 KEGG orthologs - KOs per sample) were assigned using the KEGG database to identify changes in bacterial functions and metabolic pathways associated with MS duration. In concordance with the changes detected at the taxonomic level, no significant differences in beta-diversity were found between MS individuals and controls using the KO abundance data (PERMANOVA, *p* = 0,295, R2 = 0.023). However, a higher difference between groups was detected when MS patients were stratified by disease length (**Fig. 2A**, PERMANOVA, *p* = 0.066, R2 = 0.064). In support of an overall shift of the functional potential of the microbiome on MS patients, a higher microbiome dissimilarity was found between control subjects and MS patients (both long- and short-MS) as compared to the dissimilarity detected within controls (ANOVA, p=0.0001). In concordance with the results obtained at the taxonomic level (**Fig. 1E**), the functional potential of the microbiomes of long-MS patients differed significantly more from controls (ANOVA & FDR, q<0.0001, mean difference 0.046, **Fig. 2B**) than those from short-MS patients (ANOVA & FDR, q=0.004, mean difference 0.031, **Fig. 2B**). Analysis of the KO richness showed a similar pattern as the taxonomic richness (**Fig. 1E** vs. **Fig. 2C**). A slightly higher KO richness was detected in short-MS patients and a decrease in both richness and diversity was detected in long-MS patients, albeit the observed differences were not found to be statistically significant (ANOVA, p>0.05). Next, we explored in more detailed changes in specific KOs and metabolic pathways that were associated with MS duration. An increase in the abundance of specific KOs was mainly detected in short-MS patients, while KO depletion was the most frequently detected change in long-MS individuals (**Fig. 2D**, Mann-Whitney test, p<0.0001). Subsequently, a hypergeometric test was applied to detect metabolic pathways enriched with KOs whose abundance significantly differ among groups of individuals. As expected, short-MS and long-MS patients differ from controls in different metabolic pathways. While short-MS microbiomes were enriched in ribosome encoding genes, the fecal microbiomes of long-MS individuals were enriched with glycosyltransferases (**Fig. 2E**). On the other hand, a depletion in gene orthologs of the starch/sucrose metabolic pathway was detected in short-MS individuals, while the microbiome of long-MS patients was depleted on genes for transfer RNA biogenesis (**Fig. 2E**). In summary, MS progression was associated with changes in specific gut microbial communities which subsequently impacted the overall functional potential of the microbiome. While patients with a shorter MS disease duration had on average a higher abundance of specific KOs, patients suffering MS for a longer period of time harbor microbiomes that were in general less taxonomically rich and diverse and with lower functional potential.

**Figure 2.**
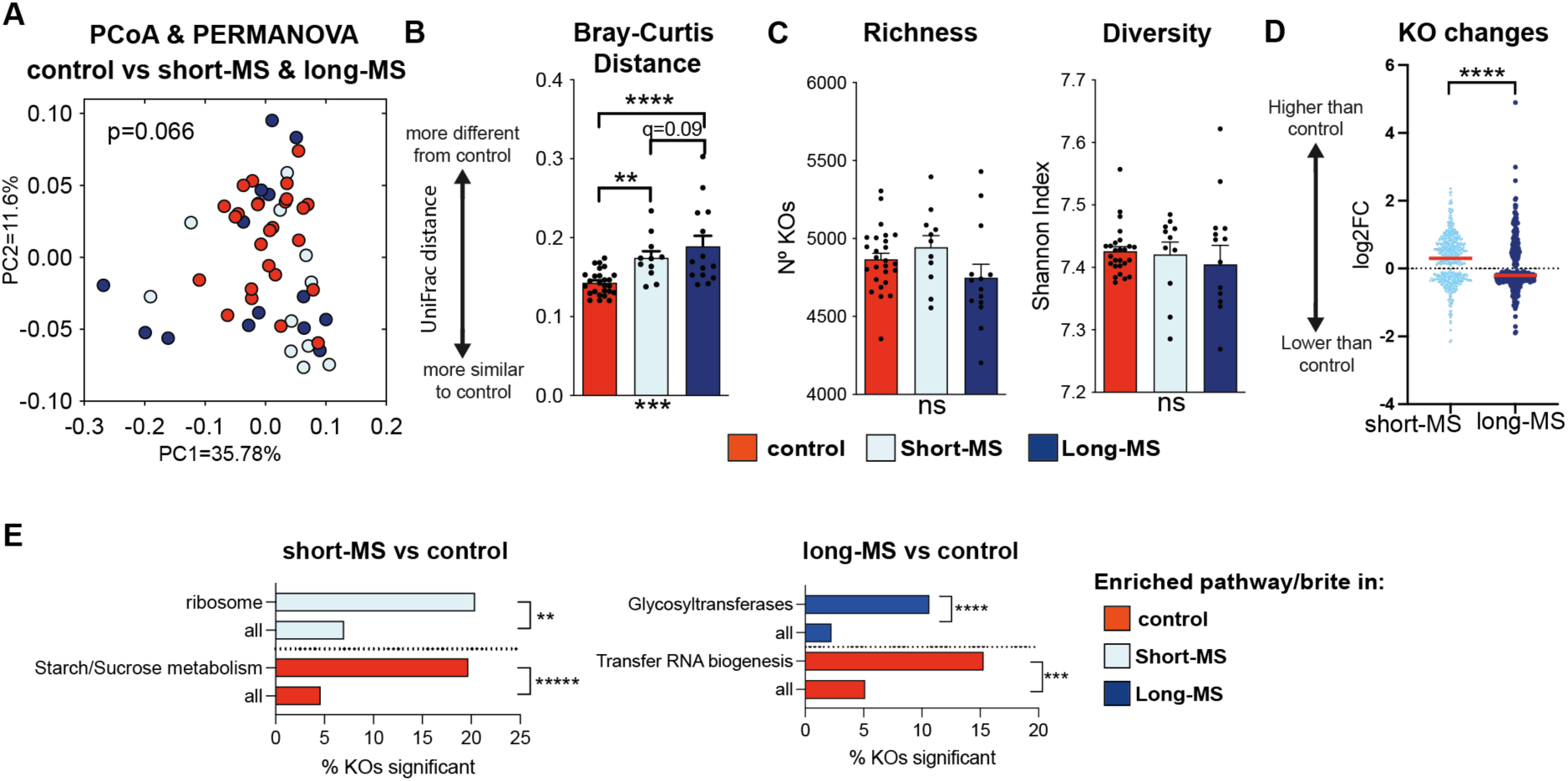
Changes in the functional microbiome potential are associated with MS duration. **(A)** PCoA and PERMANOVA analysis shows that the microbiomes (at KO level) of control and MS individuals differ when disease duration is taken into account. **(B)** Bray-Curtis distance (using KO abundance data) from controls is significantly higher to long-MS than to short-MS patients. **(C)** Similar differences in the KO richness and diversity as the ones detected at the ASV level are found in control, short-MS and long-MS individuals, although differences were not found to be statistically significant. **(D)** Log2FC between the abundance of specific KOs in short-MS vs control subjects or between long-MS vs control subjects. Each point represents one KO. Only the log2FC of KOs whose abundance is significantly different between groups is shown (p<0.05, Mann-Whitney test). On average, a significant increase in the abundance of KOs is detected in short-MS subjects, while a KO depletion is found in long-MS subjects. (**E**) Metabolic pathways significantly enrich with KOs whose abundance significantly differ between groups. *p/q<0.05, **p/q<0.01, ***p/q<0.001, ****p/q<0.0001; ANOVA and FDR in (B), ANOVA in (C), Mann-Whitney in (D), Hypergeometric test in (E). When FDR test is applied q values instead of p values are indicated. The exact q values are shown in (B) when q is between 0,05 and 0,1. Bars represent the average in (B and C) and the % of KOs within a pathway or within all pathways whose abundance is significantly increased in the group depicted in (E). Whiskers represent the SEM in (B and C). Horizontal lines represent the median in (D). N=11-26 human subjects.

### Human microbiota shows differential potential to alter CNS autoimmunity in mice

Given the differential microbiome compositions and potential functions present in MS patients, according to time from onset, we next addressed whether these changes could translate into differential disease development *in vivo*. To this end, we employed the well-established fecal microbiota transplant (FMT) model with subsequent induction of experimental autoimmune encephalomyelitis (EAE) in genetically susceptible mice that were born and raised with a fully competent microbiome (Álvarez-López et al., 2025). We and others have previously shown that the gut microbiota is a critical regulator of EAE susceptibility and severity (Berer et al., 2017; Regen et al., 2021). At the same time, FMT of human material into experimental mice has proven as valuable tool to functionally assess the influence of microbiota-derived factors on extra-intestinal pathologies (Benakis et al., 2016), including MS (Yoon et al., 2025).

To create humanized experimental animals, C57BL/6J wild type mice were treated with a broad range antibiotic cocktail to deplete the microbiota before its reconstitution with three consecutive transplants of the human donor material (**Fig. 3A**). For each human subtype (controls, short-MS and long-MS), three donors, representative of the microbial composition and disease duration of the respective cohort (**Suppl. Fig. 8**), were selected. Using the material from each of these nine human donors, three to four mice received an identical transplant in order to account for the biological variability of the FMT methodology. After a 3-week reconstitution period, we took fecal samples for microbiome analysis in order to verify that the microbiome from human donors was able to engraft in the gastrointestinal tract of recipient mice (**Fig. 3A**). To this end, we first calculated the distance at the ASV level between the microbiome of recipient mice and the microbiome of human donors. The distance at the ASV level was chosen because this is the phylotype level with higher potential for discriminating bacteria from different donors. We reasoned that, if microbial engraftment was achieved, the recipient mouse microbiome should be significantly more similar to the matching human donor microbiome than to other human donors. As shown in **Suppl. Fig.9A**, the mouse microbiomes were significantly more similar to their matching donors than to unrelated donors (*p* < 0.0001, *t*-test). Moreover, no differences in the level of engraftment were seen between groups of mice (controls, short-MS and long-MS, **Suppl. Fig 8B,** Kruskal-Wallis test, p = 0.613). Next, we analyzed if major microbiome features that distinguish the three human groups under study could also be detected in recipient mice. In concordance with the human data, the highest dissimilarity was detected between mice reconstituted with the microbiomes of long-MS patients vs. the microbiome from controls (**Suppl. Fig. 10A**, ANOVA & FDR, p<0.05). In addition, as described in humans, mice reconstituted with the long-MS patients feces contained the least diverse and least rich microbiome of all 3 groups (**Suppl. Fig. 10B**). Moreover, at the taxonomic level (from phyla to genera), the majority (86,6%) of the engrafted taxa, that significantly differ between short-MS vs long-MS individuals, followed a similar tendency on the differential abundance found in human subjects and on their respective recipient mice (**Suppl. Fig. 10C**).

**Figure 3.**
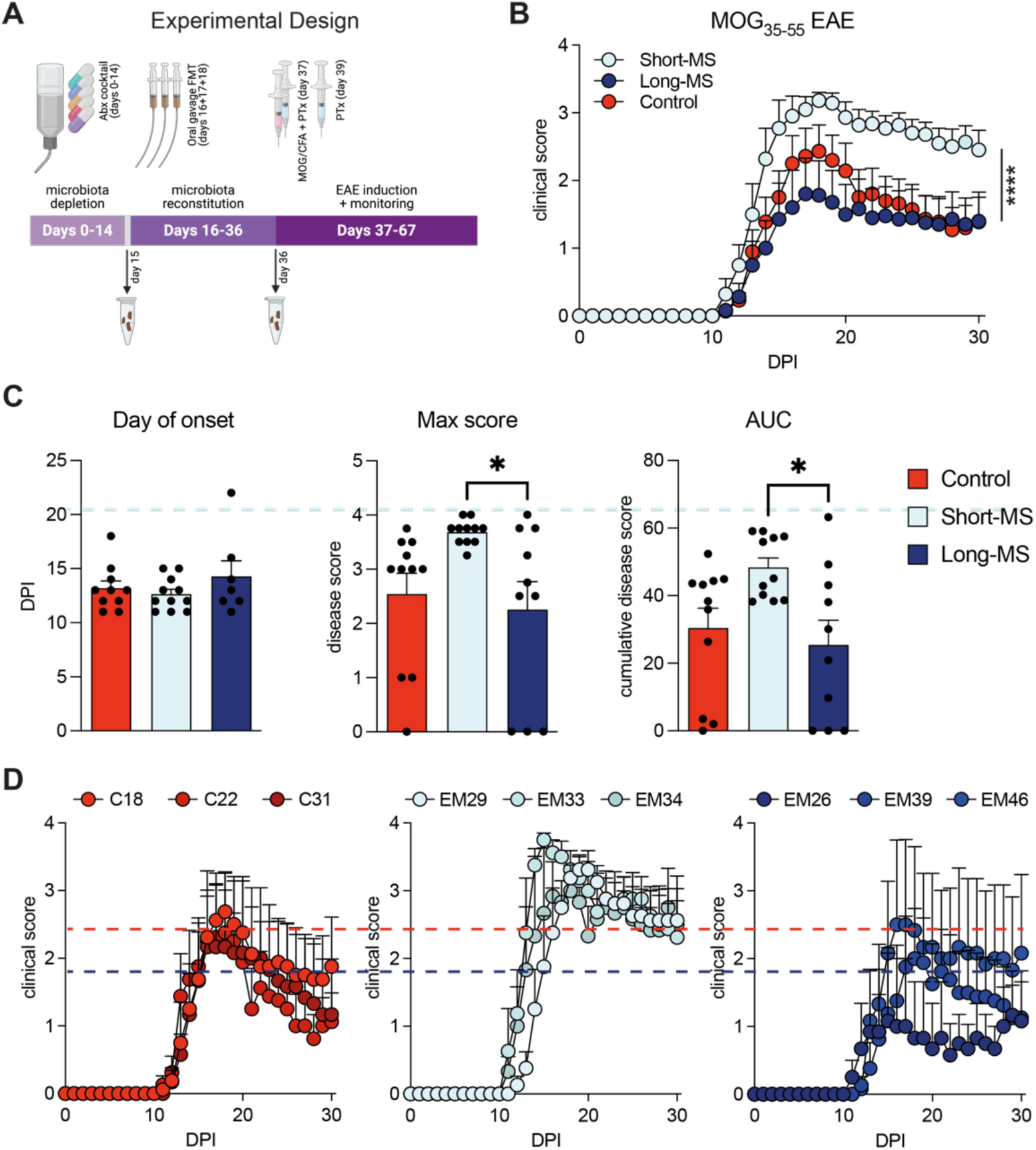
Human faecal microbiota transplants differentially alter EAE in mice. **(A)** Experimental design of the *in vivo* study. **(B)** Short-MS FMT induces significantly stronger EAE in C57BL/6 mice as compared to long-MS samples. **(C)** While all groups showed a similar day of disease onset, mice transplanted with Short-MS donor material displayed a higher disease peak (Max) score as well as higher cumulative (AUC = area under curve) disease severity, compared to the long-MS group. **(D)** Splitting the EAE clinical data from (A) into individual human donor subgroups reveals highly homogeneous responses among donors from the same groups. Alpha-numeric codes indicate the individual donor IDs. Dashed lines indicate the respective overall group disease peak according to the color code (red = Control, light blue = Short-MS, dark blue = Long-MS). *p <0.05, ****p <0.0001; 2-way ANOVA in (B), 1-way ANOVA with Tukey’s multiple comparison test in (D). Symbols in (B) represent the mean + SEM of n=10-11 animals per group. Symbols in (C) represent the mean + SEM of n=3-4 animals per group. Bars in (D) represent the mean + SEM of n=10-11 animals per group (animals that did not develop clinical symptoms were not considered for Day of onset).

Having shown that our mouse model recapitulated microbiome differences as present in our patient cohort, we next analyzed how these microbiome differences could impact disease development. We choose the EAE model as it best recapitulates the principal pathological processes of RRMS. Transplanted mice were immunized with MOG_35-55_ peptide in complete Freund’s adjuvant accompanied by two injections of Pertussis toxin (PTx), and the disease development was monitored over a period of 30 days. As can be seen in **Fig. 3B**, mice transplanted with fecal material of either controls or long-MS donors developed a similar disease course with a clearly distinguishable peak of disease followed by a remission phase. In striking contrast, mice transplanted with short-MS donor material presented with an overall higher disease burden from which they also did not recover after the peak of disease (**Fig. 3B**). This was further supported by a significantly higher cumulative disease score as well as higher disease maxima in the short-MS group, while all three groups had a similar time of disease onset (**Fig. 3C**). It is important to note that within each group (controls, short- and long-MS), the three donor sub-groups showed very homogenous EAE disease courses, thereby further supporting the validity of the overall differences detected between cohorts (**Fig. 3D)**.

EAE is driven by autoreactive CD4^+^ T cells and is in general considered to be propagated by a complex network of different immune cells. In order to estimate a potential influence of the different human microbiota transplants on the general immune compartment in FMT recipient mice, we sacrificed EAE mice at day 30 after disease induction and analyzed the spleen as major immune hub, as well as the colonic lamina propria, representing the organ rich in immune cells and physically closest to the microbiota. As can be seen in **Suppl. Fig. 11A**, the spleen displayed a generally unaltered lymphocyte composition, when comparing between the different cohorts. Likewise, the number and quality of MOG autoantigen-reactive T cells found in the spleen was not altered between groups (**Suppl. Fig. 11C-D**), pointing to a generally intact peripheral (auto)immune response. Myeloid cells are important contributors to both the pathological but also the disease-clearing mechanisms in EAE. We therefore analyzed the splenic myeloid cell compartment in high resolution, using high-dimensional flow cytometry. As can be seen in **Suppl. Fig. 12**, the major myeloid cell populations that can be identified in the spleen remained largely unaltered between cohorts. Similar to the spleen, also the colonic lamina propria remained largely unchanged when comparing the three groups. This was true for microbiota-sensitive group 1 and group 3 innate lymphoid cells (**Fig. 4A-C**), the major lymphocyte populations (**Fig. 4D**), as well as myeloid cells and their subsets (**Fig. 4E**). Together, these data argue against major changes in the immune compartment that could held responsible for the differential phenotype observed in the EAE model.

**Figure 4.**
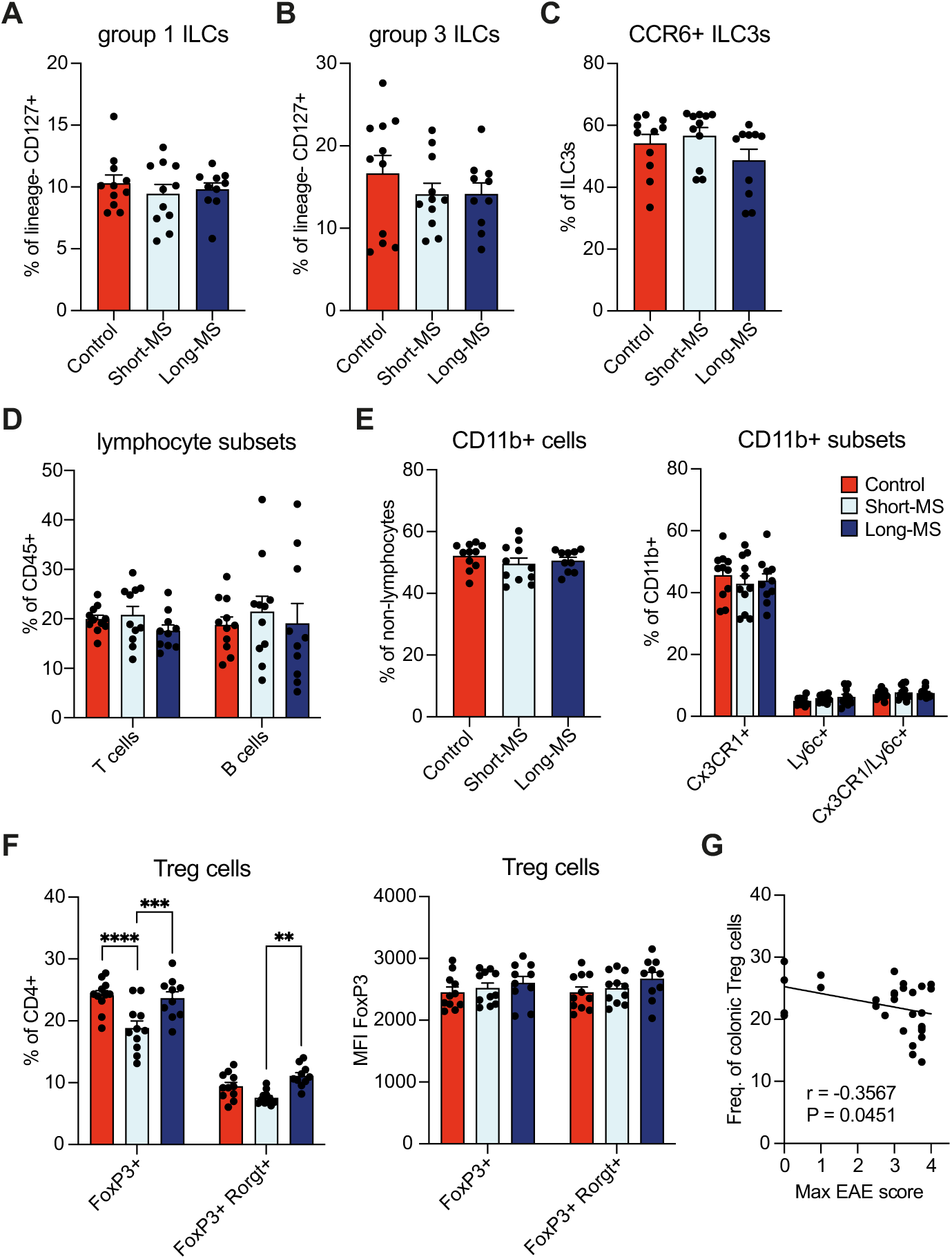
Short-MS faecal material reduces intestinal Treg cells in FMT recipient mice. **(A-C)** ILC populations in the colonic lamina propria remain unaltered regardless of the human donor group. **(D-E)** Likewise, human FMTs do not alter differentially the colonic populations of lymphocytes or CD11b^+^ myeloid cells. **(F)** Populations of large intestinal Treg cells are reduced in mice receiving short-MS human donor FMTs, while the mean fluorescence intensity (MFI) of the major Treg cell transcription factor FoxP3 remained unaltered throughout both populations in all groups. **(G)** The frequency of colonic Treg cells negatively correlates with maximal EAE severity in mice across all FMT groups. **p<0.01, ***p <0.001, ****p<0.0001, in (G) the actual p value is indicated; 2-way ANOVA with Šidák’s multiple comparison test in (F), Pearson correlation in (G). Bars in (A-F) represent the mean + SEM of n=10-11 animals per group. Line in (G) depicts the simple linear regression of the represented data points.

Ultimately, CD4^+^ T cell responses are controlled by a number of mechanisms, most importantly by regulatory T (Treg) cells, either directly or indirectly through myeloid mediators. We hypothesized that differential disease outcomes could be associated to an altered capacity of Treg cells to control autoreactive T cells. While we did not find a differential abundance of Treg cells in the spleens of EAE mice (**Suppl. Fig. 11B**), we observed a significant reduction of Treg cells in the colon of mice receiving the short-MS FMT. This reduction was found in both colonic Treg cell subsets, the thymus-derived FoxP3^+^ tTreg cells as well as the microbiota-dependent FoxP3^+^Rorgt^+^ pTreg cells (**Fig. 4F**). Notably, Treg cells from all groups presented with unchanged expression levels of FoxP3 (**Fig. 4F**). The relevance of the altered Treg frequencies was further supported by the fact that across all three cohorts, the frequency of colonic FoxP3^+^ Treg cells negatively correlated with the maximum clinical score reached during EAE development (**Fig. 4G**).

Taken together, our functional EAE data suggests that the human microbiota has a differential potential to alter the development of CNS autoimmunity in mice, depending on the MS disease duration of the human donor. Remarkably, the microbiota of short-MS patients induced more severe EAE in FMT mice, suggesting a more pro-inflammatory (or less anti-inflammatory) potential of the transplanted microbial consortia. As a potential mechanism, we identified a differential abundance of intestinal Treg cells after FMT, possibly accounting for a limited capacity to control autoreactive T cells in short-MS colonized mice (O’Connor and Anderton, 2008; Sumida et al., 2024).

### Faecal supernatants from short-MS patients induce higher pro-inflammatory cytokine secretion by human peripheral blood mononuclear cells (PBMC)

Besides Treg cell differentiation in the colonic lamina propria, which can impact CNS inflammation, microbiota-derived products can also reach the bloodstream and affect secretion of cytokines by different immune cell types (Vinolo et al., 2011; Li et al., 2018; Essex et al., 2024). Considering this possibility, we tested the ability of faecal microbiome supernatants, derived from our three human cohorts, to differentially promote the secretion of cytokines by peripheral blood mononuclear cells (PBMC). We specifically studied the secretion of TNF-α, IL-17a, IFN-γ and IL-10, since these four cytokines play a major role in the inflammatory process during MS (Valenzuela et al., 2007; Kwilasz et al., 2015; Waisman et al., 2015; Talbot et al., 2022; Carmona et al., 2025). All faecal supernatants induced an increase in IL-10 production by PBMC. However, supernatants from controls were significantly higher IL-10 inducers than supernatants from short or long-MS patients (**Fig. 5A**). No differences were detected for IFN-γ levels after incubation with the different supernatants (data not shown). Notably, a significantly higher secretion of TNF-α and IL-17a was detected in PBMC cultures incubated exclusively with faecal supernatants from short-MS patients, as compared to naïve cells (not incubated with faecal supernatants) (**Fig. 5B-C**), whereas incubation with both controls or long-MS supernatants did not enhance the secretion of these cytokines. Altogether these results indicate that the microbiome from short-MS patients has a higher capacity of inducing a systemic inflammatory response, and accordingly a higher potential to induce EAE in the mouse model.

**Figure 5.**
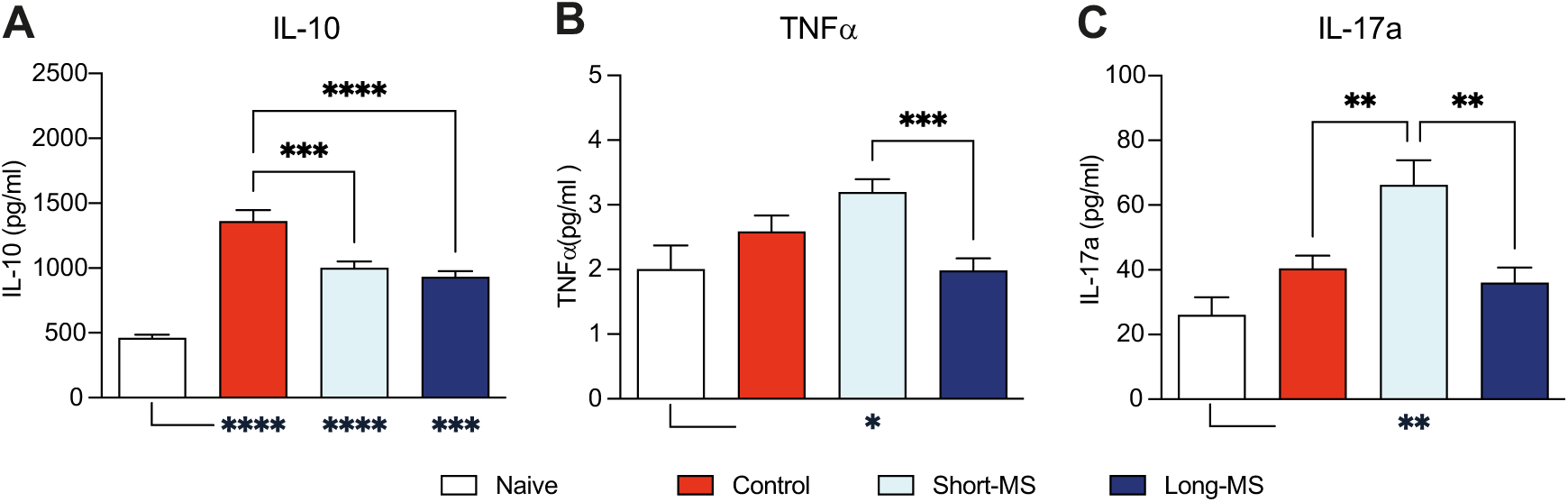
Short-MS fecal material induces proinflammatory cytokine release in human PBMCs. Cytokine concentration ((A) IL-10; (B) TNFα; (C) IL-17a) in supernatants from PBMC cultures obtained from healthy donors. Cultures were incubated with PBS (naïve), or with fecal material isolated from controls, short-MS or long-MS patients. *p <0.05, **p<0.01, ***p <0.001, ****p<0.0001; 1-way ANOVA with Tukey’s multiple comparison test in (A-C). Bars represent the mean + SEM of n=16-48 cultures in (A), n=12-35 cultures in (B) and n=7-24 cultures in (C). Asterisk symbols below the x-axis depict the statistical results comparing the Naïve group to HLT, short-MS and long-MS, respectively.

## DISCUSSION

The aetiology of MS is still a matter of debate. However, there is evidence that both genetic and environmental risk factors contribute to the development and progression of MS (Sawcer et al., 2011; Scazzone et al., 2021). In recent years, microbiome alterations have been reported in MS patients (Miyake et al., 2015; Jangi et al., 2016; Tremlett et al., 2016; Horton et al., 2021). These findings, together with the observed disease exacerbation in EAE mice after faecal transplantation from MS patients (Cekanaviciute et al., 2017), suggest that the microbiome is an important environmental risk factor in MS.

Previous studies focused on comparing microbiome composition between MS and healthy controls, between MS subtypes (RRMS vs PMS) and also between MS patients under different therapeutic regimens (Zhou et al., 2022). Here, we report, for the first time, differences in the microbiome composition within the same MS subtype (RRMS) associated with the time course of the disease. Multiple factors could alter the gut microbiome composition during the course of the disease and contribute to the changes detected. Indeed, we found specific variables that were associated with disease duration and that could be involved in the microbiome differences observed between short and long-RRMS patients. As expected, age was significantly higher in long-MS individuals. Multiple studies have reported changes in microbial taxa associated with aging (Bradley and Haran, 2024). However, none of the taxonomic differences detected between short- and long-MS individuals were associated with age, suggesting that age was not a major factor influencing microbiome changes in our MS cohort. In contrast, we found that depression and disease severity (EDSS score) were associated with the majority of changes that distinguished short-MS from long-MS microbiomes. In line with this result, several studies have identified differences in the gut microbiota of subjects suffering depression (Barandouzi et al., 2020). Although some of these changes may precede depression and may be involved in its progression, it has also been documented that antidepressant drugs can impact bacterial growth *in vitro* and alter the microbiome composition *in vivo* (Lukić et al., 2019; Rukavishnikov et al., 2023). Thus, some of the changes detected in patients with longer MS may be due to a higher usage of antidepressants by these subjects, which are more prone to develop depression. On the other hand, patients with higher disability, which were more prevalent in the long-MS group, contained lower levels of specific taxa. Within these taxa, we found that *Methanobrevibacter* genus and a specific ASV of this genus were depleted in patients with higher EDSS score. *Methanobrevibacter* is a strictly anaerobe archaea that produces methane, a molecule with neuroprotective properties, including the suppression of microglia and astrocyte activation and the reduction of inflammatory cell infiltration (Poles et al., 2019). Thus, a depletion of this genus may favour neurodegeneration and MS disability. Nevertheless, this genus was found in another study to be enriched in untreated MS individuals as compared to healthy controls (Mirza et al., 2022), and in other inflammatory disorders like periodontitis (Horz and Conrads, 2011). Thus, its role in MS, both at the early and late phases of the disease remains to be clarified.

Since the microbiome composition is key for inducing differential immune responses, we wondered whether the microbiome changes associated with disease length could differentially impact inflammation and consequently the development of CNS autoimmunity. To estimate the inflammatory capacity of the microbiome samples under study, we used the well-established EAE model in which the immunostimulatory role of the gut microbiome has been already suggested. We show that gut microbiome-depleted mice reconstituted with the microbiome from short-MS subjects developed significantly more severe EAE as compared to mice reconstituted with healthy individual microbiota, which corroborates previous findings demonstrating the inflammatory potential of MS patient microbiota (Berer et al., 2017; Cekanaviciute et al., 2017). What was less expected was our finding that mice receiving the microbiota of long-MS individuals developed EAE that was similar, with a trend to be even milder, compared to the healthy control group. Immunophenotypic analysis of the EAE mice thereby revealed an overall unchanged immune cell composition in both the spleen, as major immune hub in EAE pathogenesis, as well as in the colonic lamina propria as the direct response sight to the microbiota transplants, with one important exception: the colonic Treg cell population was significantly reduced in short-MS mice, whereas Treg cell numbers recovered in the long-MS group, thereby mirroring the overall EAE disease phenotypes. Treg cells and their role in controlling inflammation and dampening disease progression have been well established in both EAE and MS (O’Connor and Anderton, 2008; Sumida et al., 2024). In accordance with this relation, the frequency of Treg cells in the colonic lamina propria negatively correlated with the maximum disease severity across all experimental groups of mice studied here. It is therefore conceivable that the intestine is a key regulator of the systemic inflammatory response in MS and accordingly should be considered as interventional target site. In which way the gut microbiota is involved precisely in these immunological processes will have to be the subject of future studies.

To further validate the differential pro-inflammatory potential of the microbiome from different donor groups, we incubated human PBMCs (from healthy volunteers) with supernatants from the three groups of patient-derived faecal samples. We found that supernatants from short-MS patients induced higher levels of the proinflammatory cytokines IL-17a and TNF-α, whereas long-MS samples induced those cytokines in amounts comparable to the control group. TNF-α is a pleiotropic cytokine that is involved in a plethora of pathological but also physiological processes. Although it has been clearly demonstrated to take part in the pathogenesis of MS, its therapeutical targeting has remained to be challenging due to its multifaceted nature (Caminero et al., 2011). On the other hand, IL-17a is the signature cytokine of proinflammatory Th17 cells, which are well-known players in autoimmune pathologies, including MS (Waisman et al., 2015). In the context of T cell activation, both of these cytokines can therefore be regarded as proinflammatory mediators. Accordingly, our findings revealed that short-MS microbiota supernatants induced a robust inflammatory PBMC response, while the long-MS samples failed to do so.

Together, our results from the pre-clinical EAE mouse model and the *in vitro* studies with human PBMCs suggest that the microbiome differences detected in patients according to disease duration can be translated to a functionally different potential to influence systemic inflammation and consequently MS development. This inflammatory potential appears to be highest in the early phase of RRMS and declines with increasing disease duration.

One important question that remains to be clarified based on the results presented in our work is the role of microbiome changes on the progression of MS. Since later stages of MS are defined by a more brain-localized inflammation and lower systemic inflammation, it is tempting to speculate that a microbiome switch towards a less proinflammatory one may be required for progression from one MS stage to another. Conversely, it is also possible that the progression of the disease and reduction of systemic inflammation precedes and drive changes in the microbiome towards a less pro-inflammatory one. In fact, in a previous study we demonstrated that deficiency in IL-17a signaling alters the microbiome in mice (Regen et al., 2021). Most importantly, the microbiome from IL-17 KO mice was less capable of inducing inflammation in the EAE model. Thus, future studies will be important to define the role of microbiome changes on disease progression, including the identification of the specific bacterial commensals and functions involved, and elucidate the role of host immunity on microbiome compositional changes during the course of the disease.

One potential limitation of our study is that samples were not collected repeatedly from the same patients at different time points, precluding our ability to study changes in the microbiome within the same individual. Thus, follow-up longitudinal studies will have to demonstrate whether these findings can be reproduced in individual patients with sequential probing and analysis of the gut microbiota, including its functional testing. In addition, we acknowledge that our cohort was limited in size and all patients were derived from a single center. Nevertheless, we were able to validate important aspects of our findings, as based on the stratification of RRMS patients by disease length, employing data from a larger multi-center MS microbiome study (Zhou et al., 2022).

Despite these limitations, our results bring novel findings to the MS field that may have therapeutical implications. We have found that the inflammatory potential of the microbiome is higher at earlier stages of RRMS, when systemic inflammation theoretically plays a major role and when disease modifying therapies that directly target systemic inflammation are more effective (Lassmann, 2017). This implies that the microbiome may have a key role in inducing inflammation at an early stage of the disease, creating a window of opportunity for microbiome-based therapies aimed at reducing systemic inflammation and preventing MS progression. In this sense, several clinical trials have analyzed the impact of probiotic administration on MS progression showing promising results (Tankou et al., 2017, 2018; Salami et al., 2019; Rahimlou et al., 2022) Considering our results showing that short-MS patients contain a more pro-inflammatory microbiome, it is conceivable that this type of patients will benefit most from microbiome-based therapeutic approaches such as probiotics, while this type of therapies will have a limited impact on long-MS individuals containing a microbiome with lower inflammatory potential and EAE-inducing capacity. Thus, it will be important for future clinical trials to stratify patients according to disease length in order to evaluate the efficacy of microbiome-based therapies to decrease MS symptoms and define the window in which a probiotic may be effective for preventing MS progression. Moreover, our study has started to pinpoint those microbiome differences associated with disease course. The identified microbiome features, if validated, could serve as biomarkers that could facilitate patient stratification for clinical-decision making.

## MATERIALS AND METHODS

### Participant enrolment and sample collection

Twenty-seven MS patients were enrolled at Centro Hospitalar São João MS outpatient neurology clinic (Porto, Portugal). Two of these patients developed SPMS and were excluded for subsequent analysis. Among the 25 RRMS patients remaining, 11 showed a disease lower than 16 years and were included in the short-MS group; 14 patients showed a disease duration of 16 years or higher and were classified long-MS. Twenty-six controls not developing MS were enrolled among patient spouses or family-unrelated subjects, accompanying the patient at the time of the clinical appointment. The study protocol conformed to the ethical guidelines of the 1975 Declaration and ethical approval was granted (Approval ref: 250.16). Informed consent was obtained in accordance with the institutional review board regulations.

Participant information (e.g., gender, age, disease duration, MS subtype, disease severity indexes and therapy) is included in Table 1. Faecal samples were collected by each participant in appropriate provided, sterile containers and divided in aliquots for microbiota analysis, faecal transplant in mice and faecal supernatant preparation.

For establishment of primary cultures from peripheral blood mononuclear cells (PBMC), 5 healthy volunteers (3 females and 2 males, mean age 37, age interval 27-52) were enrolled among the health personnel of Centro Hospitalar São João MS outpatient neurology clinic.

### Faecal DNA extraction

Bacterial DNA from faecal samples was extracted using the QIAamp® DNA Fast Stool Mini kit (QIAGEN) as previously described (Isaac et al., 2022). Extractions were performed according to manufacturer instructions with introduction of a previous mechanic disruption step with bead-beating. Briefly, samples (∼200 mg) were resuspended in the first buffer of the extraction kit. Subsequently the samples were shaken on a Vortex-Genie 2 equipped with a Vortex Adapters (Mobio) at maximum speed for 5 min in the presence of 500 µl of glass micro-beads (acid-washed glass beads 150-212 µm, Sigma®). Then, we followed the protocol of QIAamp® DNA Fast Stool Mini kit.

### 16S rRNA sequencing and analysis

The V3-V4 regions of the 16S rRNA gene were amplified (Kapa HiFi HotStart Ready Mix), indexed with Nextera® XT Index Kit (96 indexes, 384 samples) and sequenced as described in the manual for “16S Metagenomic Sequencing Library Preparation” of the MiSeq platform (Illumina) using MiSeq Reagent Kit V3. 16S rRNA gene sequences were denoised and processed with DADA2 v1.8 (Callahan et al., 2016), as we have previously described (Yogev et al., 2022). Briefly, prior to denoising, as recommended in the DADA2 pipeline tutorial, the 3ʹ ends of the forward and reverse pair-end sequences (35 and 75 nucleotides, respectively), which contained lower quality nucleotides, were removed from each sequence. The first nucleotides (17 in the forward read and 21 in the reverse read) were also trimmed to remove the V3-V4 primers. Sequences containing undetermined nucleotides (N) were discarded. After denoising, the forward and reverse pair-end sequences were merged together, with a minimum overlapping region of 15 bases and a maximum of 1 mismatch. DADA2 and the command removeBimeraDenovo were subsequently applied to remove chimeras. Taxonomy was assigned for each ASV using the Silva 138 database (Quast et al., 2013) as reference and the naive Bayesian classifier implemented in DADA2. Shannon index was obtained at the ASV level using the package vegan v2.5-3 and the R 3.6.0 software. Validation of the results was performed using the taxonomic and diversity microbiome data deposited in the supplementary tables from a published multicentric study (Zhou et al., 2022).

### Shotgun sequencing and analysis

Metagenomic shotgun sequencing was performed using a NextSeq platform (Illumina, USA) according to the manufacture’s recommendations in order to obtain 150bp pair-end sequences. Paired-end reads were filtered out and trimmed with PRINSEQ-lite (v.0.20.4) and the following parameters: trim_qual_right: 30; trim_qual_type: mean; trim_qual_window:2 0; min_length: 50 (Schmieder and Edwards, 2011). The overlapping pairs were merged (joined) by means of FLASH (version 1.2.11) (Magoč and Salzberg, 2011) to obtain longer reads. The merged and non-merged forward reads were concatenated in a single FASTQ file and subsequently converted into FASTA format. The reads were mapped onto the *Homo sapiens* genome by means of Bowtie2 (version 2.3.4.3) (Langmead and Salzberg, 2012), with the very-sensitive preset. The aligned reads were discarded. The preprocessed reads were assembled to generate contigs with Megahit (version 1.14) (Li et al., 2016). The preprocessed reads were mapped onto the generated contigs with BLASTn (version 2.8.1+) (Camacho et al., 2009), with the following parameters: task: megablast; perc_identity: 97; qcov_hsp_perc: 90; strand: both; max_target_seqs: 1; max_hsps: 1. The unaligned reads were considered as (singleton) contigs too and added to the assembled contigs. Open reading frames (ORFs) inside (singleton and assembled) contigs were predicted with Prodigal (version 2.6.3) (Hyatt et al., 2010), with meta parameter. Overlaps between genes and read alignments coordinates were tallied by means of R (version 3.6.0) to set genes abundance (R Core Team (2019). R: A language and environment for statistical computing. R. Foundation for Statistical Computing, Vienna, Austria. URL https://www.R-project.org/). The abundance was also normalized by gene length. The predicted ORFs were annotated by means of HMMER-hmmsearch (version 3.2.1) (Eddy, 2011), maximum E-value of 1e-3, with the KEGG Orthology (KO) database as reference.

### Selection of faecal samples for animal studies and supernatant preparation

In order to select representative samples of the microbiome changes detected in the different groups of subjects (i.e. control, short-MS and long-MS), we took into account the distribution of UniFrac distance to the control group shown in 1E. We selected those samples that were closest to the median within each group and for which sufficient faecal material was available (**Suppl. Fig. 8**).

### Preparation of the human fecal transplants

For the mice FMT experiment, selected human donor samples were aliquoted and individual aliquots thawed at the respective days of transfer. For that, the frozen donor material was transferred into degassed PBS preheated to 37°C at a concentration of 1 ml per 0.5 g of stool. The samples were then vortexed at full speed for 1 min and the resulting solution quick-spun to pellet any particulate matter. The resulting supernatant was taken up into syringes that were pre-assembled with gavage needles to ensure that no particles potentially clogging the needles were transferred.

### *In vivo* mouse model

C57BL/6J male mice were purchased from Envigo/Inotiv. For intestinal microbiota depletion, mice were treated with 0.5 mg/ml Ampicillin (Ratiopharm), 0.5 mg/ml Gentamycin, 0.5 mg/ml Metronidazole, 0.5 mg/ml Neomycin and 0.25 mg/ml Vancomycin, all dissolved in autoclaved drinking water, for 14 days. The antibiotics cocktail was freshly prepared every 3 days. At day 15, mice were put back on autoclaved drinking water without antibiotics and during a 24 h wash-out period. At day 16, the human donor transplants were freshly prepared as described above, and 200 µl immediately transferred to antibiotic-treated recipient mice by oral gavage. This procedure was repeated 24 h (day 17) and 48 h (day 18) later. For each human donor sample, 3-4 mice were reconstituted. The transplants were left to engraft for a total of 21 days, after which a faecal sample was collected to control for the engraftment efficiency. Immediately after the engraftment period, EAE was induced. All animal experimentation was approved by the local administration and was performed in accordance with federal and state policies.

### EAE induction

Active EAE was induced as previously described (Regen et al., 2021). Briefly, mice were immunized with 50 µg MOG35-55 peptide (GenScript) emulsified in complete Freund’s Adjuvant (CFA, BD Biosciences) supplemented with 10 mg/ml of heat-inactivated Mycobacterium tuberculosis H37RA (BD Biosciences). The emulsion was injected s.c. at the tail base. Mice also received 200 ng pertussis toxin (List Biological Laboratories) as i.p. injection at the day of immunization as well as 2 days later. Clinical assessment of EAE was performed daily according to the following criteria: 0, no disease; 1, decreased tail tone; 2, abnormal gait (ataxia) and/or impaired righting reflex (hind limb weakness or partial paralysis); 3, partial hind limb paralysis; 3.5, complete hind limb paralysis; 4, hind limb paralysis with partial fore limp paralysis; 5, complete fore limp paralysis or moribund and 6, dead.

### *Ex vivo* flow cytometry analysis

After the 30-day clinical observation period, EAE mice were humanely sacrificed and organs collected. Single cell solutions from spleen and colon were prepared as previously described (Regen et al., 2021; Shanmugavadivu et al., 2024). Cells were stained for flow cytometry in 96-well plates. Fc receptors were blocked by incubation with anti-mouse CD16/32 Fc blocking solution for 20 minutes at 4°C. After washing, cells were incubated for 30 minutes at 4°C in the dark in 50 μL of a cocktail containing surface-binding antibodies and a viability dye in FACS buffer, followed by washing. Cells were either acquired directly on a FACSCantoII (BD Biosciences), FACSymphony A3 (BD Biosciences) for multidimensional myeloid cell analysis, or fixed overnight with 2% formaldehyde and acquired within one week. For intranuclear staining of the transcription factor Foxp3 as a marker for regulatory T cells, cells were stained for surface markers and then fixed, permeabilized and stained using the Foxp3 transcription factor kit (eBioscience) according to manufacturer’s recommendations. For the MOG antigen recall assay, spleen cells were cultured in the presence of 20 µg/ml MOG peptide. After 2 h, cultures were supplemented with 1 µg/ml Brefeldin A (Merck) in order to block the secretion of cytokines. After additional 4 h of culture, cells were harvested and stained for flow cytometry analysis. After surface staining, cells were fixed and permeabilized with the Cytofix/Cytoperm kit (BD Biosciences) according to manufacturer’s recommendations and stained for intracellular cytokines. T cell expression of CD154 was assessed as marker for recent activation, thereby serving as surrogate marker for MOG antigen specificity. CD154high T cells were then further analysed for cytokine expression. All flow cytometry data was analysed using FlowJo software. For dimensionality reduction analysis cells were pre-gated to exclude doublets, dead cells, and lymphocytes (CD19^+^ B cells and CD3^+^ T cells). CD11b^+^ events were randomly downsampled using the Downsample plugin and samples from each condition were concatenated into one FCS file. Dimensionality reduction was performed using the t-SNE plugin. The FlowSOM plugin was used to cluster cells and ClusterExplorer plugin was used to investigate marker expression for manual cluster annotation.

### Faecal supernatant preparation

Faecal supernatants were prepared from an aliquot of selected faecal samples, as described above. The protocol used was an adaptation from previous reports (Meelu et al., 2014; He et al., 2019). Aliquots were thawed on ice and resuspended in PBS (Sigma-Aldrich, Merck) at a concentration of 1 g/5 ml. Suspensions were then centrifuged at 2550g at 4°C for 20 minutes and supernatant was collected and filtered through a 40 μm cell strainer. The filtered suspension was centrifuged again at 13000g at 4°C for 15 minutes. The resulting supernatant was collected, filtered through a 0,45 μm filter (agarose acetate membrane; Whatman, Merck) and filtered again through a 0,22 μm filter. The resulting suspension was divided in aliquots and stored at −80°C until addition to cell cultures.

### Human PBMC isolation and culture

PBMC were isolated from healthy donors using Histopaque-1077^TM^-based cell centrifugation as described before (Nazareth et al., 2015). After collection and washing of mononuclear cells, cells were resuspended in RPMI 1640 with GlutaMAX^TM^ (Invitrogen) supplemented with antibiotic/antimycotic solution (Invitrogen) and 10% FBS (complete RPMI). Cells were then seeded on flat-bottom 96-well plates at a density of 5×10^4^ cells/ml (200 μL/well), incubated with the faecal supernatants selected from MS patients and HLT controls as described and cultured for 24h at 37°C and 5% CO_2_.

### Cytokine determination

For cytokine quantification on PBMC culture supernatants, four experiments were set up in different days using cells isolated from each of four healthy donors. After cell isolation and 24h incubation with 0.5 and 1% of the selected faecal supernatants, total PBMC culture supernatant from each well (200 μL) was collected and divided in four 50 μL aliquots, which were frozen at −20°C. Determination of TNF-α, IL-17, IFN-γ and IL-10 was performed using specific human Proquantum^TM^ Immunoassay Kits (Thermo Fisher Scientific), which are designed to detect target proteins using amplification by qPCR. Each aliquot was tested in duplicate using 2 μL of sample, and read on an Applied Biosystems^TM^ real time PCR instrument.

### Statistical analysis

To analyze global differences in the microbiota composition, the unweighted UniFrac distances between pairs of samples were calculated using the phangorn_2.4.0 and GUniFrac_1.1 packages for tree construction using ASV sequencing data and computation of UniFrac distance, respectively. Subsequently, PCoA analysis was performed using the calculated UniFrac distances, the package vegan v2.5-3 and the R 3.4.0 software. In order to analyze community-level differences in the microbiome among groups of samples, a non-parametric test, permutational multivariate analysis of variance (PERMANOVA), was applied using the adonis function from the R vegan package.

To identify bacterial taxa that could discriminate between groups of samples, the Boruta algorithm was applied (Kursa and Ruknicki, 2010), as we have previously described (Artacho et al., 2024). Subsequently, analysis of compositions of microbiomes with bias correction (ANCOM-BC) was applied (Lin and Peddada, 2020) in order to validate those features identified and confirmed by Boruta. Only prevalent taxa were included in this analysis (i.e. those taxa present in at least 60% of samples within at least one of the groups under comparison and having a relative abundance five times higher than the minimum relative abundance detected in all samples). To adjust for multiple hypothesis testing and obtaining q values, we used the FDR approach by Benjamini and Hochberg (Benjamini and Hockberg, 1995), implemented in the p.adjust function of the R stats package v.3.6.0. Differences with a p value lower than 0.05 and q value lower than 0.1 were considered significant.

The non-parametric Mann-Whitney test was also applied for the identification of significant differences in the abundance of KOs among groups of samples. A hypergeometric test and FDR approach was subsequently applied in order to identify metabolic pathways that were enriched with KOs whose abundance significantly increase or decrease after transplant (p<0.05). A pathway was considered significant if more than 10 genes within the pathway were found to increased or decreased comparing a pair of groups and the hypergeometric q value was lower than 0.1.

The Spearman correlation test was applied to identify associations between continuous variables and taxonomic relative abundances.

The Shapiro test was applied in order to evaluate if continuous variables such as diversity, richness, UniFrac distance or cytokine levels followed a normal distribution. If all groups under comparison followed a normal distribution, a parametric test was applied, otherwise, a non-parametric test was used. For two-groups comparisons, either the parametric T-test or the non-parametric Mann-Whitney test were applied. For more than two-groups comparison, the parametric one-way ANOVA test or the non-parametric Kruskall-Wallis test with False Discovery Rate for multiple comparisons were applied. The Fisher test was applied for categorical variables analysis.

## ACKNOWLEDGEMENTS

The authors would like to thank Michaela Blanfeld and David C. Uhlfelder for their assistance with experimental procedures and data analysis. This work was funded by a grant of Merck s.a., by Associação para a Investigação do Líquido Cefalorraquidiano of Hospital São João, by the Deutsche Forschungsgemeinschaft (DFG, German Research Foundation) under project number 490846870 –TRR355/1 to AW and TR, and project number 532695030 to TR, and by the Spanish MICINN [PID2023-150086OB-I00] to CU.

## FIGURES

**Supplementary Figure 1.**
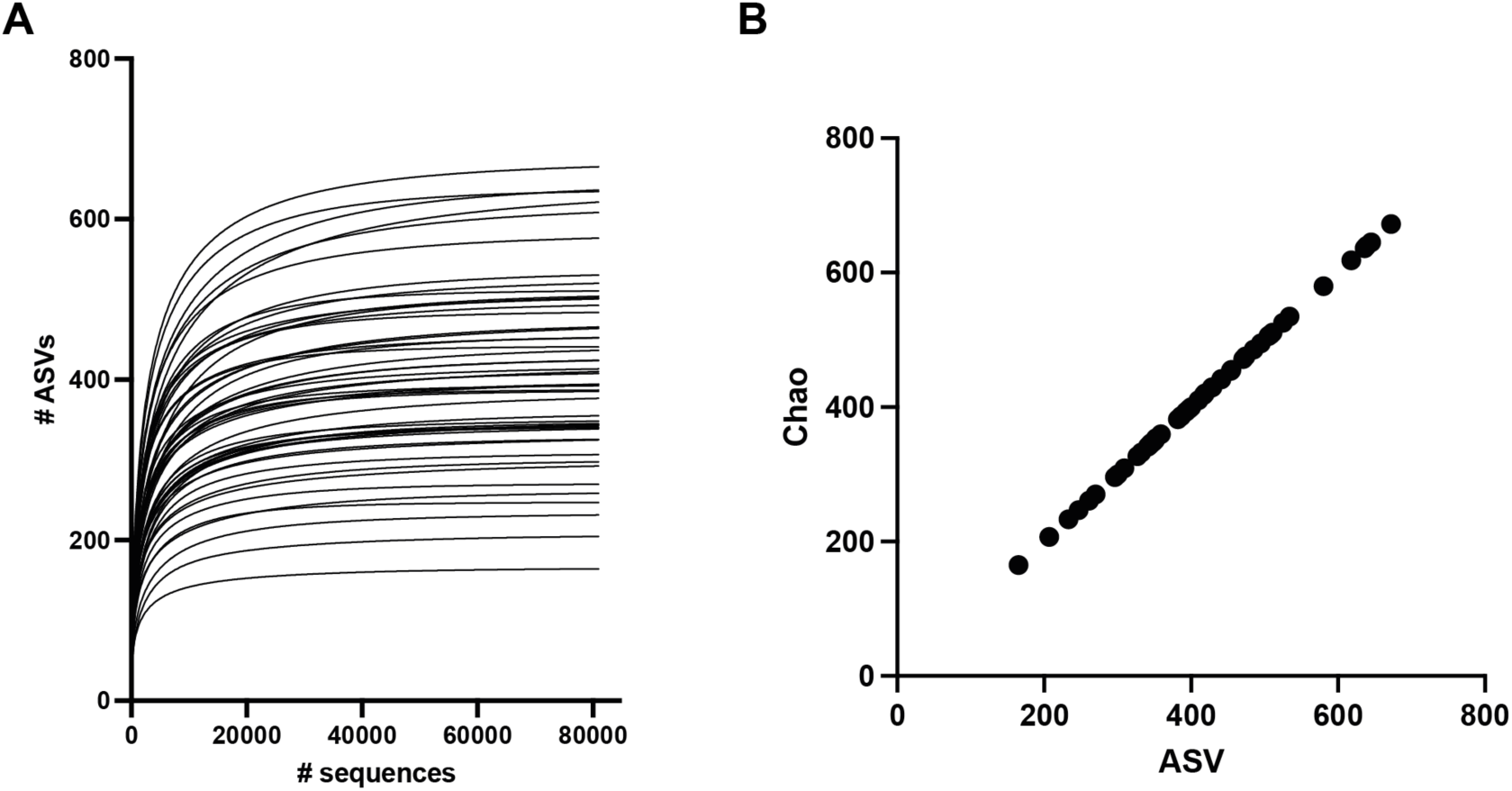
16s rRNA gene sequencing depth was sufficient to identify the majority of ASVs. **(A)** Rarefaction curves representing the number of ASVs identified in each sample after analyzing an increasing number of sequences. The rarefaction curve was done with a maximum of 81000 sequences per sample (minimum number of sequences found in a sample). **(B)** The number of ASVs is almost identical to the Chao index, representing the potential maximum number of ASVs that could be detected. N=51 samples.

**Supplementary Figure 2.**
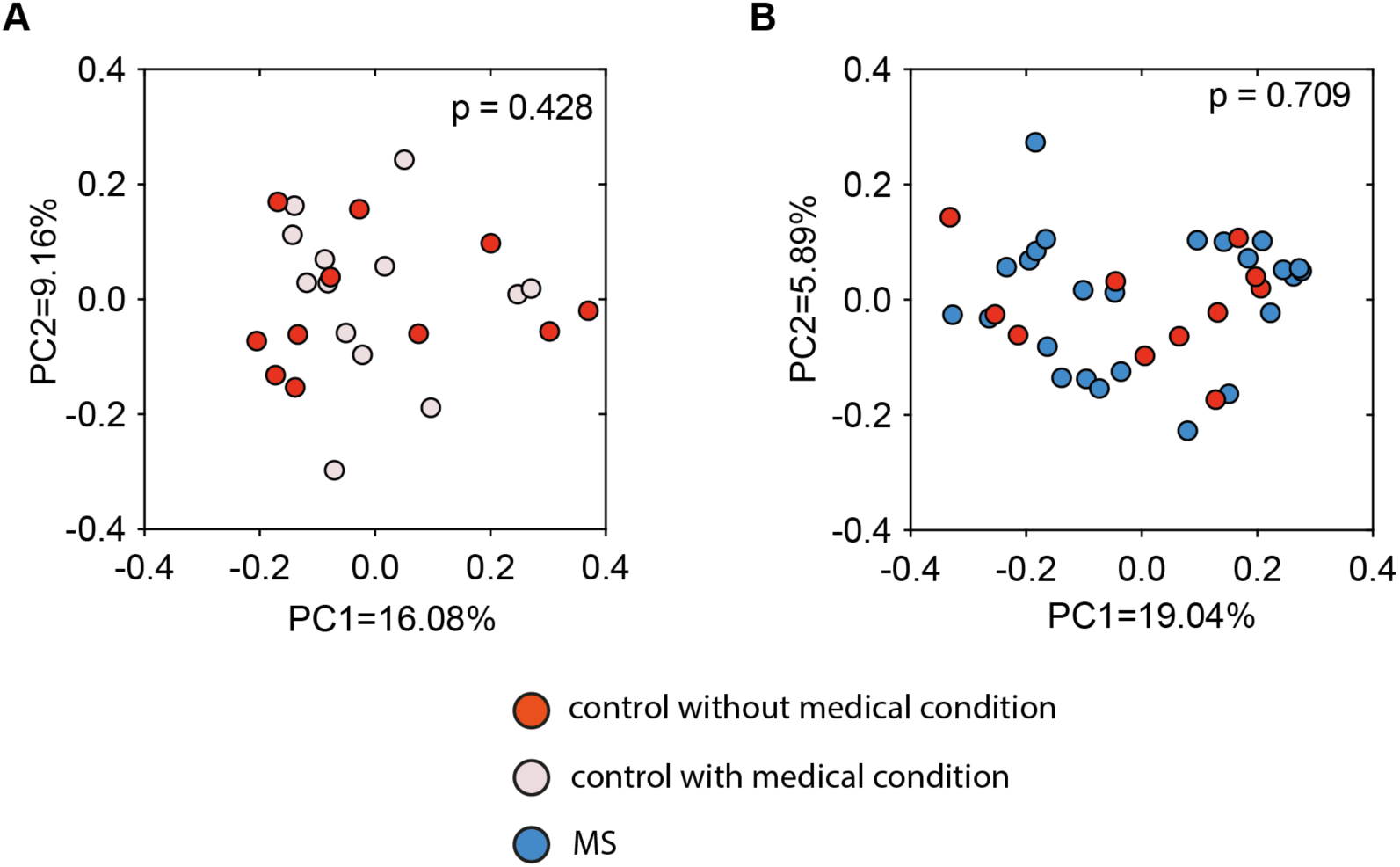
Medical conditions of control individuals do not impact microbiome composition. **(A)** PCoA using unweighted UniFrac distances between samples from controls with or without medical conditions. **(B)** PCoA using unweighted UniFrac distances between samples from controls without medical conditions and MS patients. P values indicated in the figures were calculated using the PERMANOVA test.

**Supplementary Figure 3.**
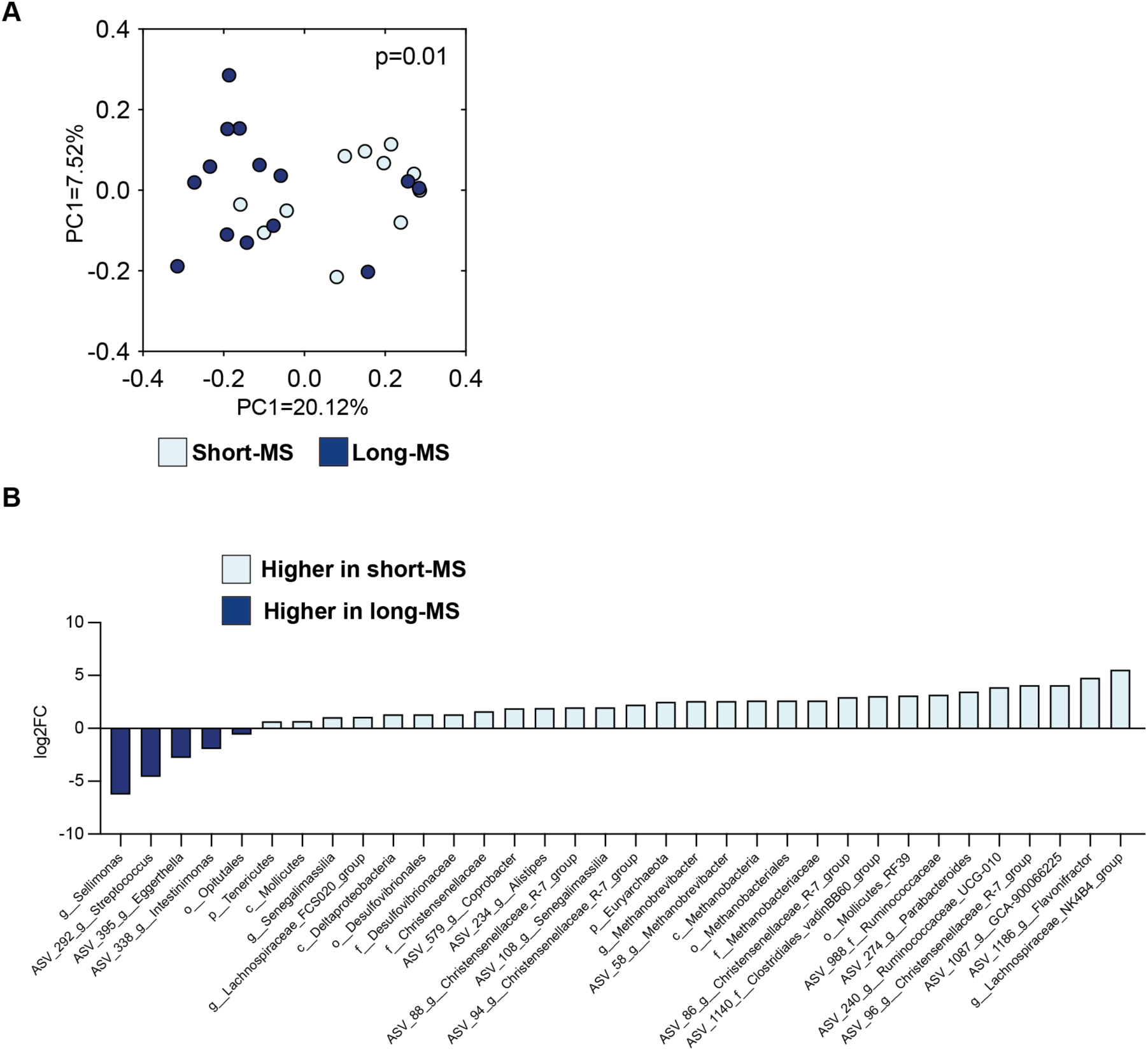
Disease duration is associated with differences in the microbiome in MS patients. **(A)** PCoA using unweighted UniFrac distances between samples from MS patients which were divided into groups depending if they had a disease duration ≥ 16 years (the median disease duration) or not. The P value indicated in the figure was calculated using the PERMANOVA test. **(B)** Significant differences at different taxonomic levels (phylum-ASV) found between short-MS patients and long-MS patients. Boruta and ANCOM-BC (p<0.05). Bars represent the log2FC of the relative abundance between both groups of patients.

**Supplementary Figure 4.**
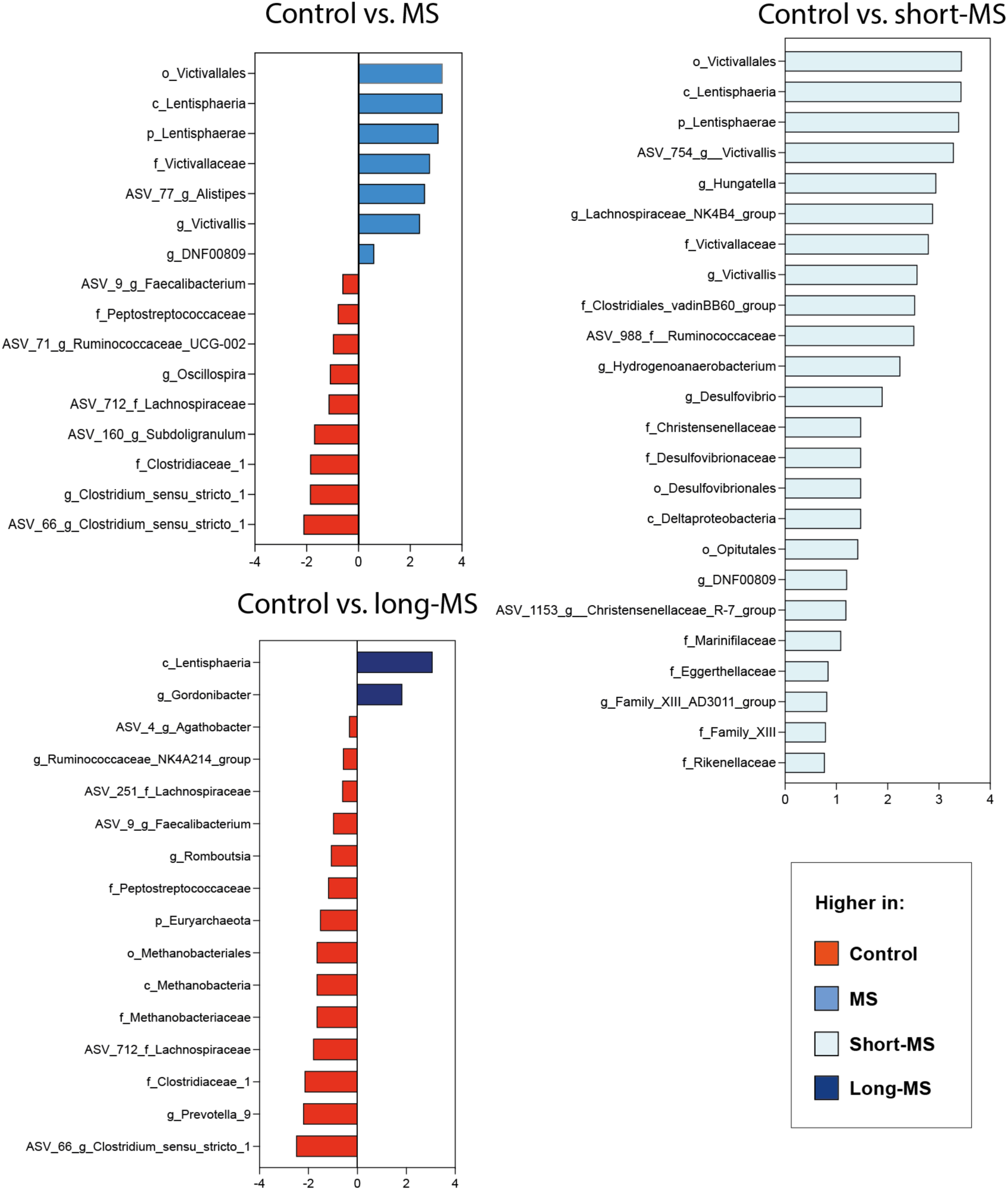
Differences in microbial taxa between controls and MS patients. Significant differences at different taxonomic levels (phylum-ASV) found between different groups of patients as indicated. Boruta and ANCOM-BC (p<0.05). Bars represent the log2FC of the relative abundance between both groups of patients.

**Supplementary Figure 5.**
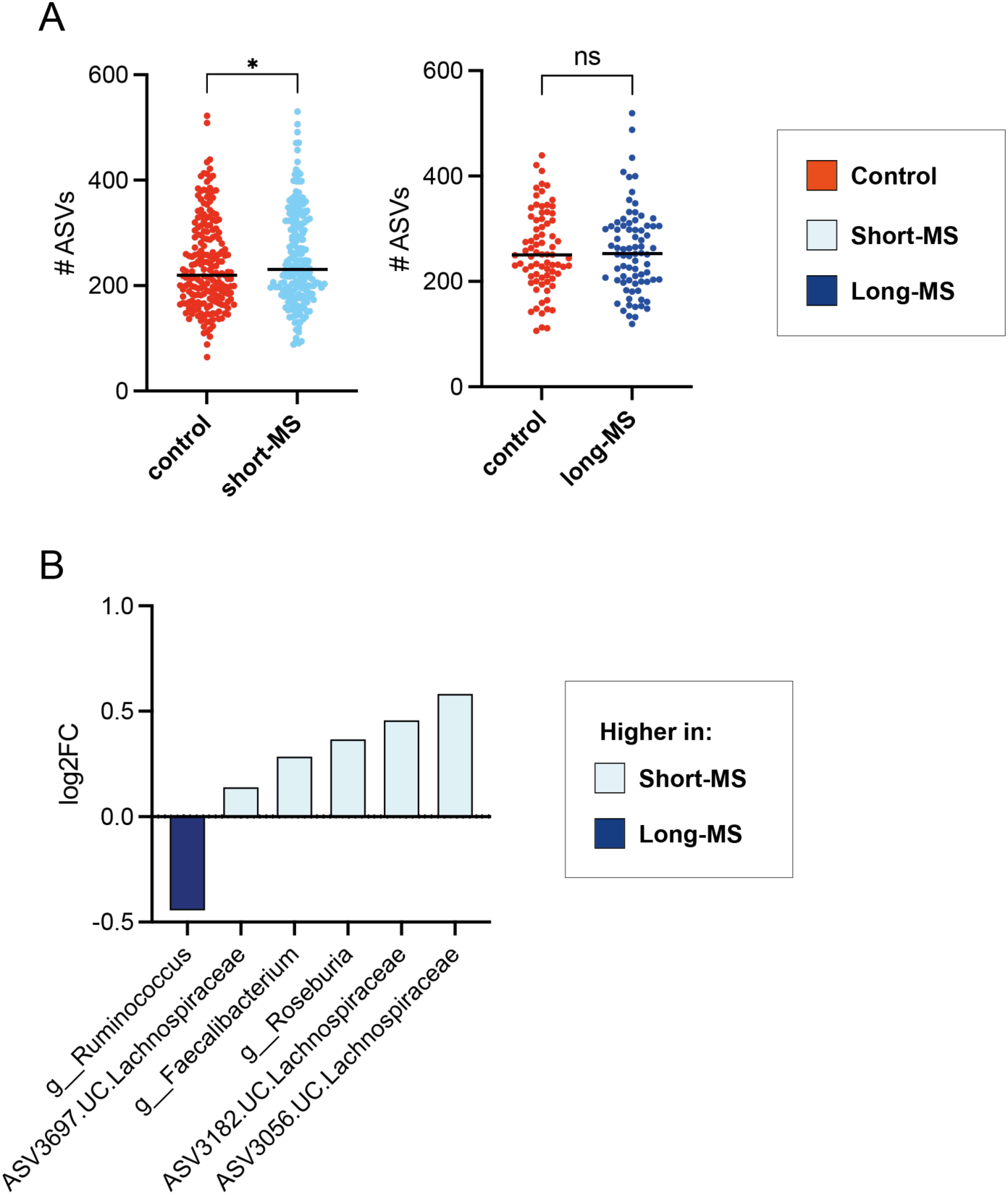
Differences in the microbiome detected between short-MS and long-MS patients in a larger cohort. (**A**) Microbiome richness, measured as the number of ASVs detected, was significantly higher in short-MS patients as compared to house-hold controls (Wilcoxon matched-pairs signed rank test, *p <* 0.05). (**B**) Bacterial genera and ASVs with discriminatory power (Boruta algorithm) found to be significantly differ in abundance between short-MS and long-MS individuals (Ancom-BC, *p <* 0.05). N= 207 short-MS and 78 long-MS.

**Supplementary Figure 6.**
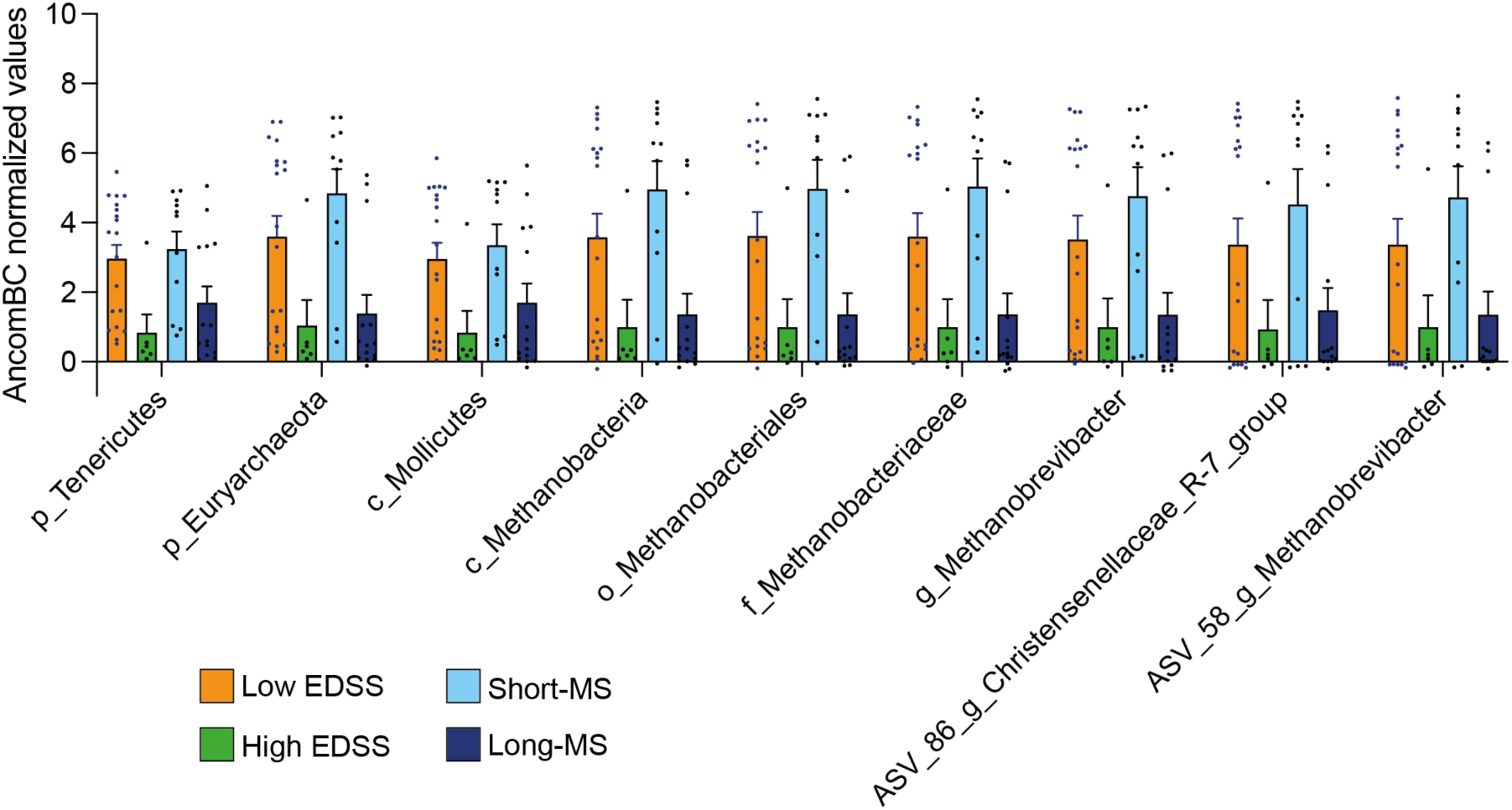
Taxa differentially abundant between short- and long-MS patients that are significantly associated with EDSS score. Taxa whose abundance is significantly different between MS patients with low EDSS (≤ 3) and high EDSS (>3). ANCOM-BC p<0.05. Bars represent the median of ANCOMBC normalized values of these two groups of patients. For comparison, the values for short-MS and long-MS groups were also included in the figure. Only taxa that were found to be significantly different between short-MS and long-MS patients (Suppl. Fig. 3B) were studied.

**Supplementary Figure 7.**
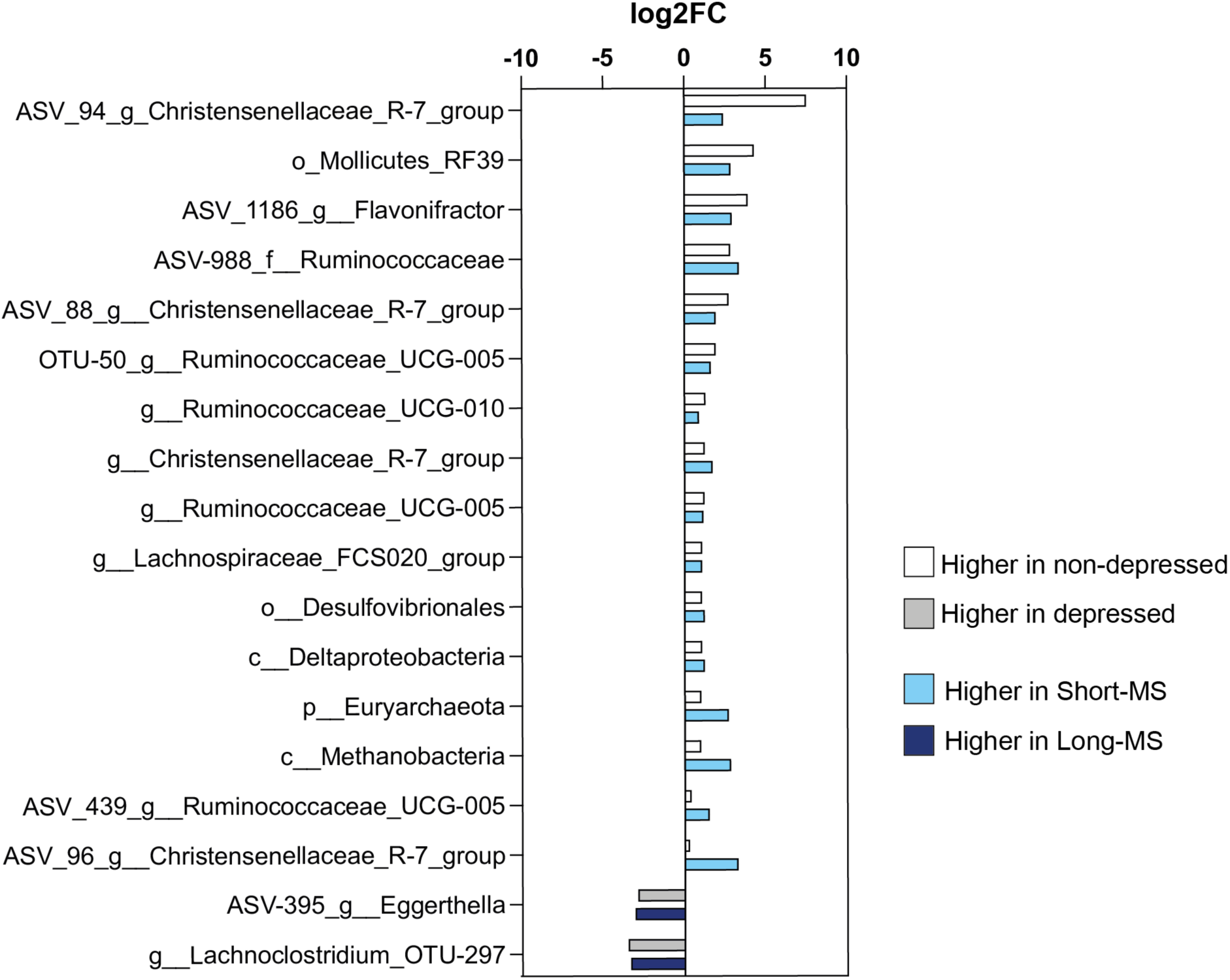
Taxa differentially abundant between short- and long-MS patients that are significantly associated with depression. Taxa whose abundance is significantly different between MS patients with or without depression. ANCOM-BC p<0.05. Bars represent the log2FC of the relative abundance of the indicated taxa between the two groups under comparison. For comparison, the log2FC of the relative abundance of the same taxa between short-MS and long-MS patients is shown.

**Supplementary Figure 8.**
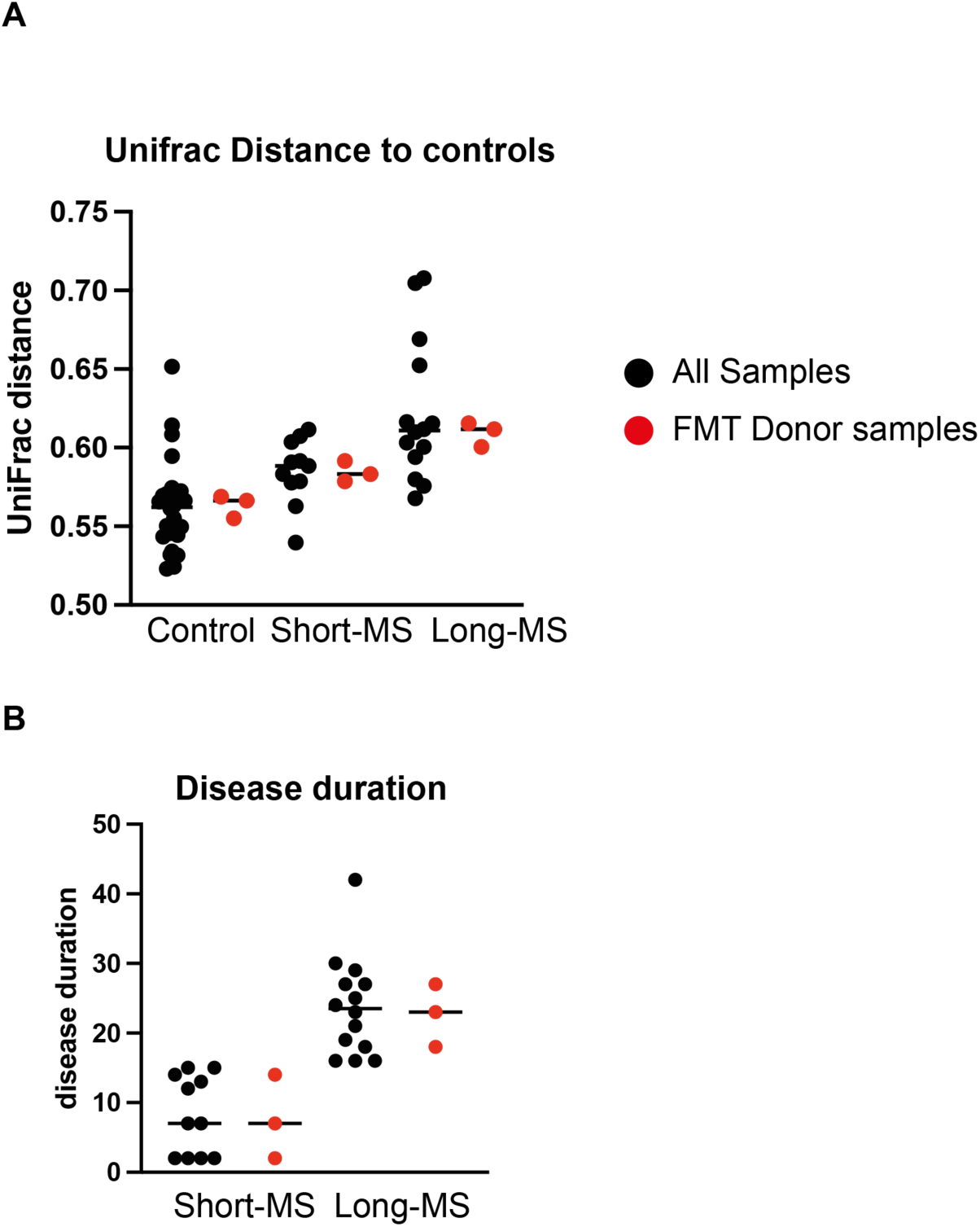
Selection of donor samples for performing FMT in mice. **(A)** Unifrac distance to control samples. **(B)** Disease duration of MS patients. In black are indicated all individuals within a group. In red are indicated those that were chosen as donors for the FMT mouse experiments. Samples were chosen based on proximity to the median Unifrac distance within each group, representativity of disease duration within short-long-MS groups and sample availability.

**Supplementary Figure 9.**
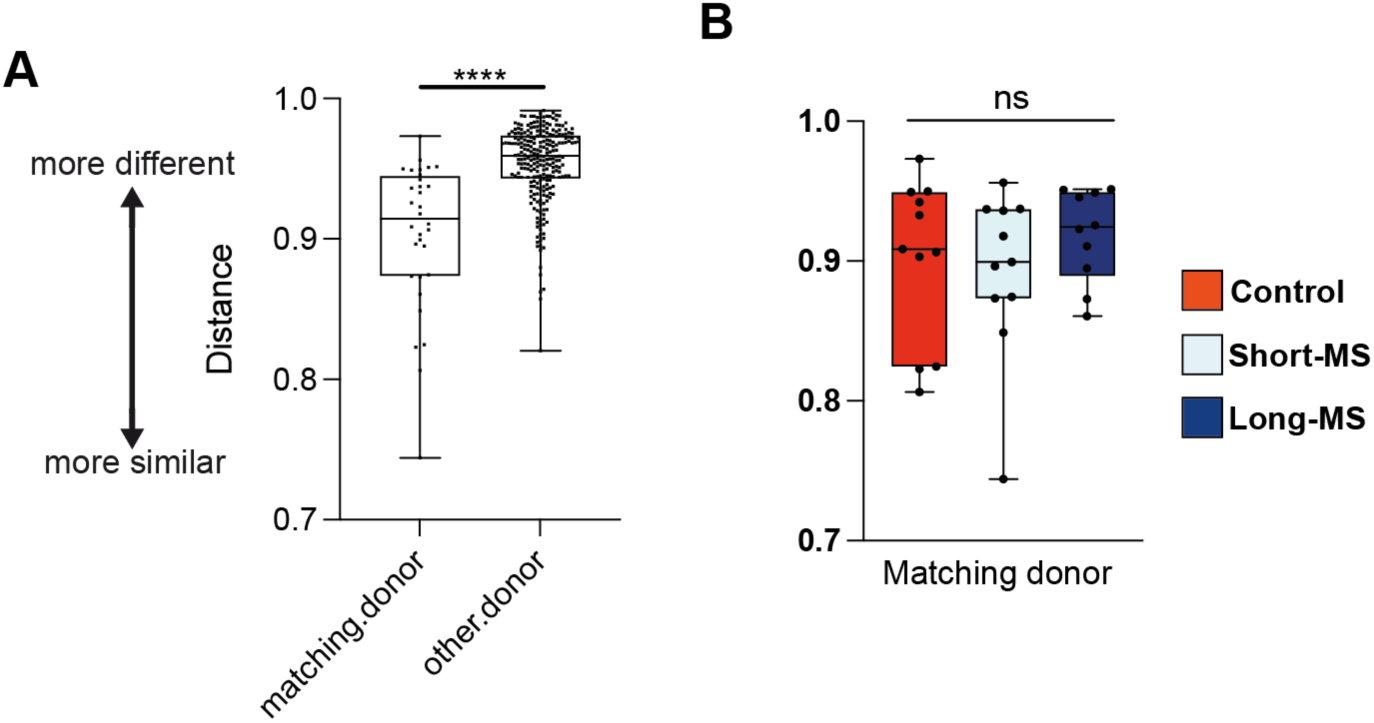
The microbiome from transplanted mice is more similar to the donor matching sample. **(A)** Bray-curtis distances between the microbiota of recipient mice and matching donors (Donor) or unrelated donors (Other). **(B)** Bray-curtis distances between recipient mice and matching donors. ****p<0.0001, T-test for A and ANOVA for B.

**Supplementary Figure 10.**
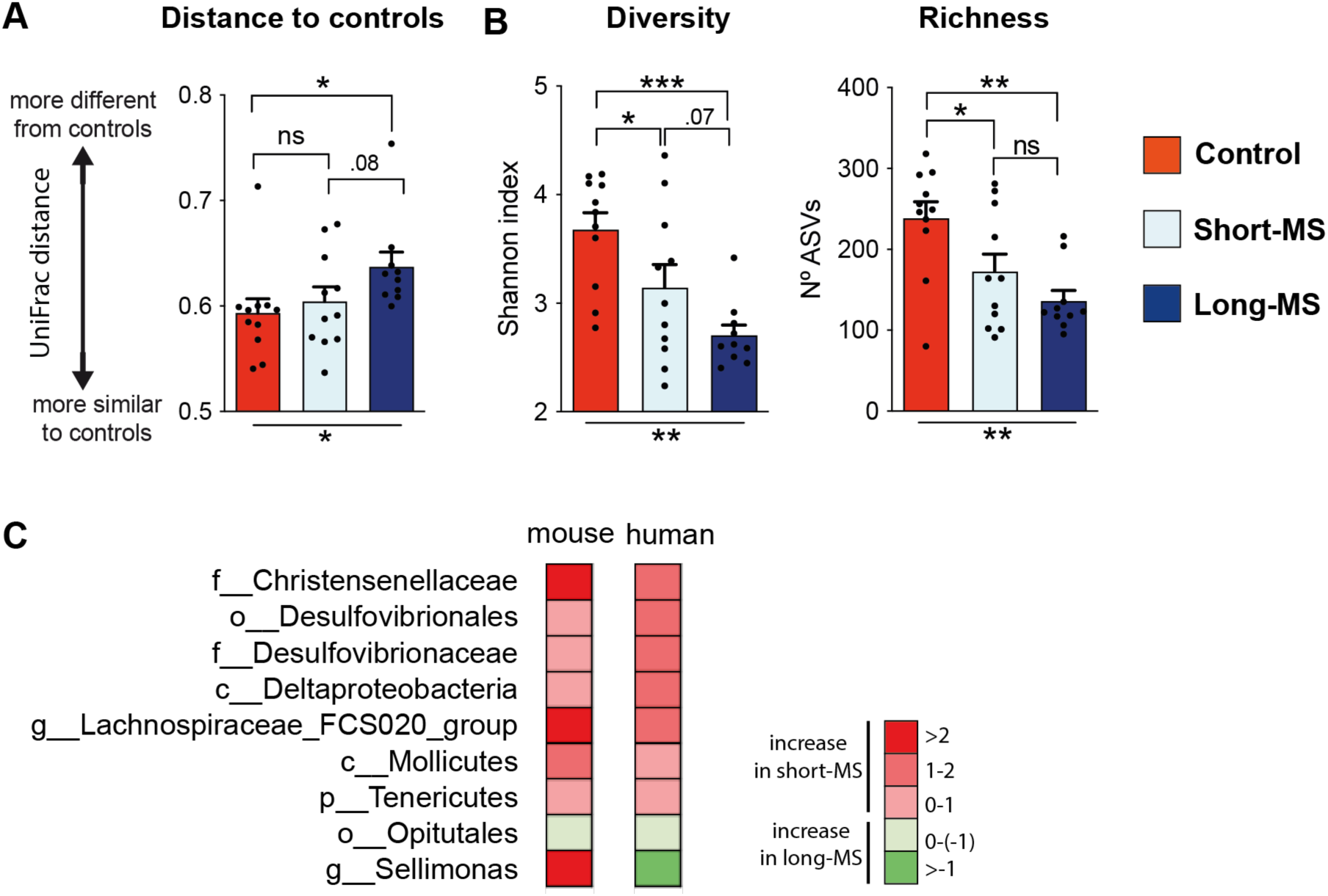
Similar microbiome differences are detected in MS groups and their respective recipient mice. **(A)** UniFrac distance from control microbiome recipient mice is significantly higher to long-MS than to short-MS microbiome recipient mice. (**B**) Richness and diversity significantly differ among mice recipient of short-MS, long-MS and control microbiomes. *p<0.05, **p<0.01, ***p<0.001, Kruskall-Wallis & FDR for (A), ANOVA & FDR for (B). Bars represent the average; whiskers represent the SEM. N=10-11 mice. **(C)** Heatmap showing the log2FC in the relative abundance of specific taxa between short-MS and long-MS patients (humans) and recipient mice reconstituted with microbiomes from short-MS or long-MS individuals. Red colors indicate higher abundances in short-MS while green colors indicate higher abundances in long-MS. Only taxa at phylum-genus level that were found to be significantly different between short-MS and long-MS patients (Suppl. Fig. 3B) were studied. ASVs were not studied because of the inter-individual variability at the ASV level and the fact that only 3 human subjects were used as donors in the mouse experiments. Only taxa that were found to engraft in at least one group of mice are shown.

**Supplementary Figure 11.**
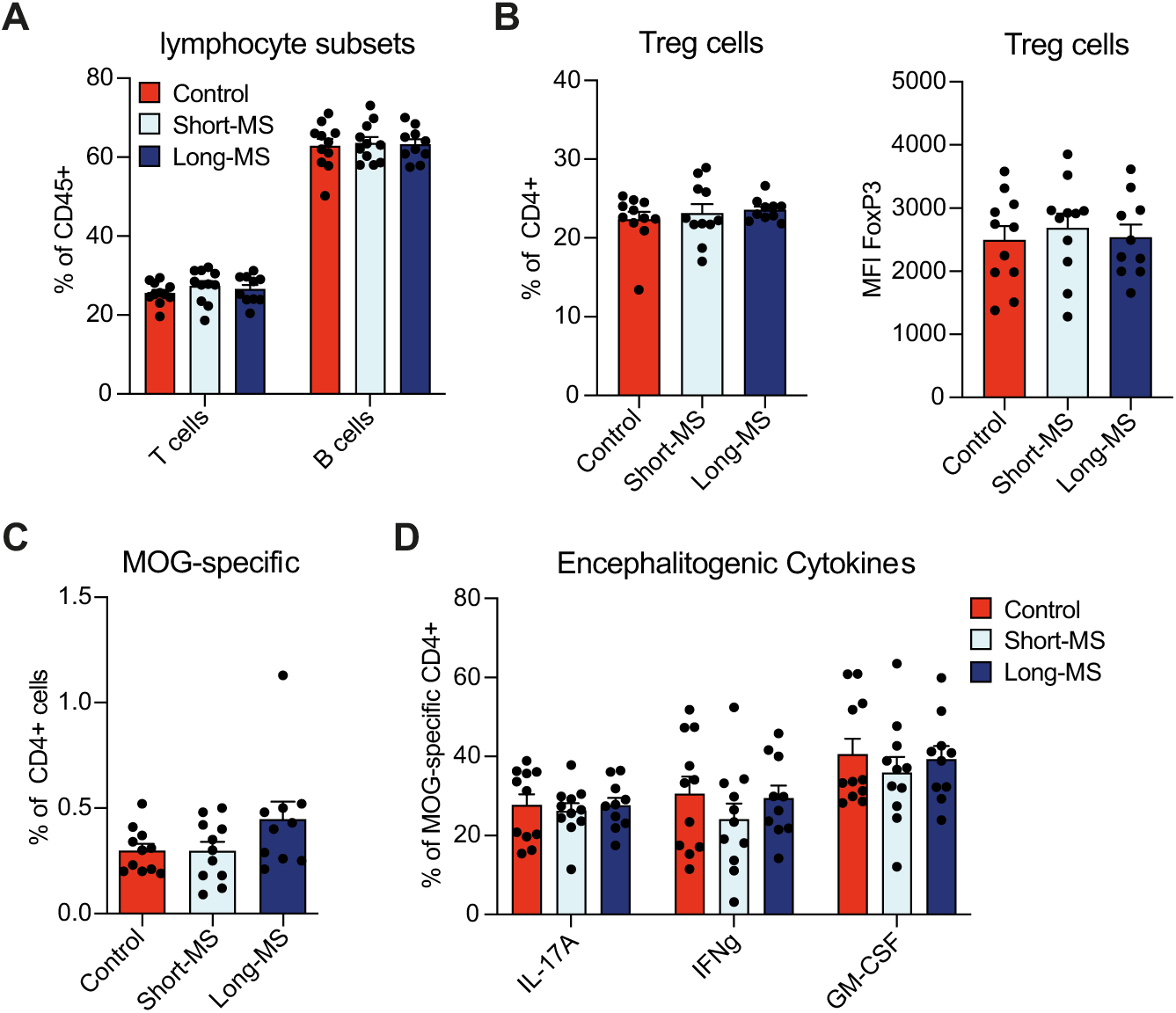
Human FMTs do not alter the splenic immune compartment and response. **(A)** The major lymphocyte populations in the spleen remain unaltered regardless of the human donor group. **(B)** Similarly, human FMTs do not alter differentially the frequency or FoxP3 transcription factor expression of Treg cells in the spleen. **(C-D)** A MOG antigen recall assay using splenocytes isolated at DPI30 of EAE revealed no differences in the frequency of MOG-reactive CD4^+^ T cells (C) as well as their production of EAE-relevant cytokines (D). Bars in (A-D) represent the mean + SEM of n=10-11 animals per group.

**Suppl. Fig. 12.**
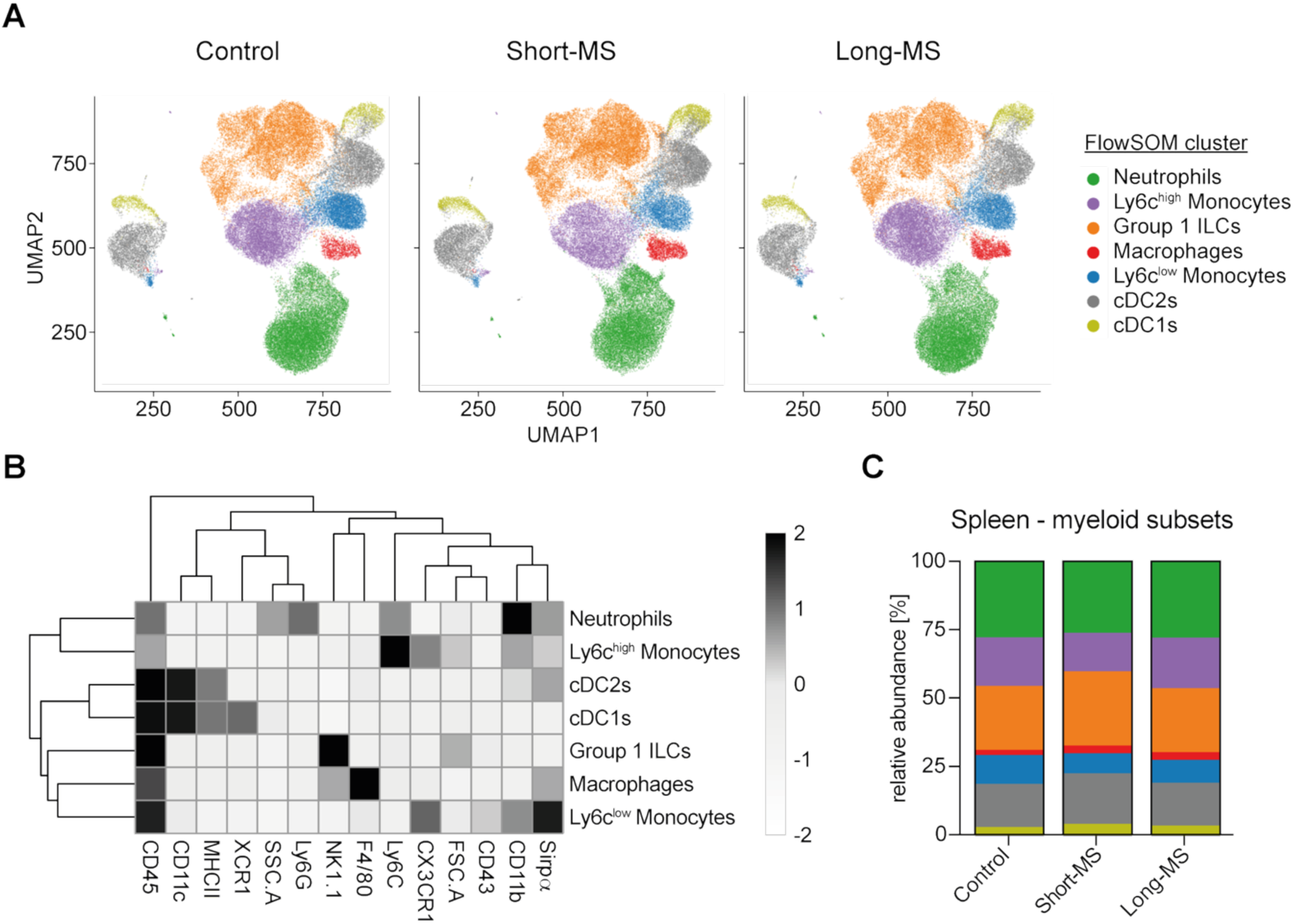
Human FMTs do not alter the splenic myeloid cell compartment. **(A-C)** Spleens of EAE mice at DPI30 were analyzed for the composition of myeloid cells by high-dimensional flow cytometry. **(A)** UMAP visualization of grouped samples after down-sampling and FlowSOM clustering. **(B)** Heat map of specific marker expression as basis for the clustering in (A). **(C)** Relative abundance and comparison between groups of the clusters as shown in (A).

## REFERENCES

1. Álvarez-López, A.I., Ponce-España, E., Cruz-Chamorro, I., Santos-Sánchez, G., Bejarano, I., Álvarez-Sánchez, N., Lardone, P.J., Carrillo-Vico, A., 2025. Differences of the 6N and 6J Substrains of C57BL/6 Mice in the Development of Experimental Autoimmune Encephalomyelitis. MedComm (2020) 6, e70228. doi:10.1002/mco2.70228

2. Artacho, A., González-Torres, C., Gómez-Cebrián, N., Moles-Poveda, P., Pons, J., Jiménez, N., Casanova, M. J., Montoro, J., Balaguer, A., Villalba, M., Chorão, P., Puchades-Carrasco, L., Sanz, J. & Ubeda, C. Multimodal analysis identifies microbiome changes linked to stem cell transplantation-associated diseases. Microbiome 12, 229 (2024).

3. Barandouzi, Z.A., Starkweather, A.R., Henderson, W.A., Gyamfi, A., Cong, X.S., 2020. Altered Composition of Gut Microbiota in Depression: A Systematic Review. Front. Psychiatry 11, 541. doi:10.3389/fpsyt.2020.00541

4. Baranzini, S.E., Oksenberg, J.R., 2017. The Genetics of Multiple Sclerosis: From 0 to 200 in 50 Years. Trends Genet. 33, 960–970. doi:10.1016/j.tig.2017.09.004

5. Benakis, C., Brea, D., Caballero, S., Faraco, G., Moore, J., Murphy, M., Sita, G., Racchumi, G., Ling, L., Pamer, E.G., Iadecola, C., Anrather, J., 2016. Commensal microbiota affects ischemic stroke outcome by regulating intestinal γδ T cells. Nat. Med. 22, 516–523. doi:10.1038/nm.4068

6. Benjamini, Y., Hockberg, Y., 1995. Controlling the false discovery rate: a practical and powerful approach to multiple testing. Journal of the Royal Statistical Society 57, 289–300.

7. Berer, K., Gerdes, L.A., Cekanaviciute, E., Jia, X., Xiao, L., Xia, Z., Liu, C., Klotz, L., Stauffer, U., Baranzini, S.E., Kümpfel, T., Hohlfeld, R., Krishnamoorthy, G., Wekerle, H., 2017. Gut microbiota from multiple sclerosis patients enables spontaneous autoimmune encephalomyelitis in mice. Proceedings of the National Academy of Sciences of the United States of America 114, 10719–10724. doi:10.1073/pnas.1711233114

8. Bradley, E., Haran, J., 2024. The human gut microbiome and aging. Gut Microbes 16, 2359677. doi:10.1080/19490976.2024.2359677

9. Callahan, B.J., McMurdie, P.J., Rosen, M.J., Han, A.W., Johnson, A.J.A., Holmes, S.P., 2016. DADA2: High-resolution sample inference from Illumina amplicon data. Nature Methods 13, 581–583.

10. Camacho, C., Coulouris, G., Avagyan, V., Ma, N., Papadopoulos, J., Bealer, K., Madden, T.L., 2009. BLAST+: architecture and applications. BMC Bioinform. 10, 421. doi:10.1186/1471-2105-10-421

11. Caminero, A., Comabella, M., Montalban, X., 2011. Tumor necrosis factor alpha (TNF-α), anti-TNF-α and demyelination revisited: An ongoing story. J. Neuroimmunol. 234, 1–6. doi:10.1016/j.jneuroim.2011.03.004

12. Carmona, S., Aghzadi, J., Vincent, T., Labauge, P., Carra-Dallière, C., Lehmann, S., Desplat-Jégo, S., Ayrignac, X., 2025. TWEAK and TNFɑ Pro-inflammatory Soluble Cytokines and their Specific Autoantibodies Secretion in Multiple Sclerosis Patients. Inflammation 48, 2494–2502. doi:10.1007/s10753-024-02205-0

13. Cekanaviciute, E., Yoo, B.B., Runia, T.F., Debelius, J.W., Singh, S., Nelson, C.A., Kanner, R., Bencosme, Y., Lee, Y.K., Hauser, S.L., Crabtree-Hartman, E., Sand, I.K., Gacias, M., Zhu, Y., Casaccia, P., Cree, B.A.C., Knight, R., Mazmanian, S.K., Baranzini, S.E., 2017. Gut bacteria from multiple sclerosis patients modulate human T cells and exacerbate symptoms in mouse models. Proceedings of the National Academy of Sciences of the United States of America 114, 10713–10718. doi:10.1073/pnas.1711235114

14. Cox, L.M., Maghzi, A.H., Liu, S., Tankou, S.K., Dhang, F.H., Willocq, V., Song, A., Wasén, C., Tauhid, S., Chu, R., Anderson, M.C., Jager, P.L.D., Polgar-Turcsanyi, M., Healy, B.C., Glanz, B.I., Bakshi, R., Chitnis, T., Weiner, H.L., 2021. Gut Microbiome in Progressive Multiple Sclerosis. Ann. Neurol. 89, 1195–1211. doi:10.1002/ana.26084

15. Eddy, S.R., 2011. Accelerated Profile HMM Searches. PLoS Comput. Biol. 7, e1002195. doi:10.1371/journal.pcbi.1002195

16. Essex, M., Pascual-Leone, B.M., Löber, U., Kuhring, M., Zhang, B., Brüning, U., Fritsche-Guenther, R., Krzanowski, M., Vernengo, F.F., Brumhard, S., Röwekamp, I., Bielecka, A.A., Lesker, T.R., Wyler, E., Landthaler, M., Mantei, A., Meisel, C., Caesar, S., Thibeault, C., Corman, V.M., Marko, L., Suttorp, N., Strowig, T., Kurth, F., Sander, L.E., Li, Y., Kirwan, J.A., Forslund, S.K., Opitz, B., 2024. Gut microbiota dysbiosis is associated with altered tryptophan metabolism and dysregulated inflammatory response in COVID-19. npj Biofilms Microbiomes 10, 66. doi:10.1038/s41522-024-00538-0

17. Fagnani, C., Neale, M.C., Nisticò, L., Stazi, M.A., Ricigliano, V.A., Buscarinu, M.C., Salvetti, M., Ristori, G., 2014. Twin studies in multiple sclerosis: A meta-estimation of heritability and environmentality. Mult. Scler. J. 21, 1404–1413. doi:10.1177/1352458514564492

18. Goris, A., Vandebergh, M., McCauley, J.L., Saarela, J., Cotsapas, C., 2022. Genetics of multiple sclerosis: lessons from polygenicity. Lancet Neurol. 21, 830–842. doi:10.1016/s1474-4422(22)00255-1

19. He, Y., Li, X., Yu, H., Ge, Y., Liu, Y., Qin, X., Jiang, M., Wang, X., 2019. The Functional Role of Fecal Microbiota Transplantation on Dextran Sulfate Sodium-Induced Colitis in Mice. Frontiers in Cellular and Infection microbiology 9, 393. doi:10.3389/fcimb.2019.00393

20. Horton, M.K., McCauley, K., Fadrosh, D., Fujimura, K., Graves, J., Ness, J., Wheeler, Y., Gorman, M.P., Benson, L.A., Weinstock-Guttman, B., Waldman, A., Rodriguez, M., Tillema, J., Krupp, L., Belman, A., Mar, S., Rensel, M., Chitnis, T., Casper, T.C., Rose, J., Hart, J., Shao, X., Tremlett, H., Lynch, S.V., Barcellos, L.F., Waubant, E., Centers, the U.S.N. of P.M., 2021. Gut microbiome is associated with multiple sclerosis activity in children. Ann. Clin. Transl. Neurol. 8, 1867–1883. doi:10.1002/acn3.51441

21. Horz, H.-P., Conrads, G., 2011. Methanogenic Archaea and oral infections – ways to unravel the black box. J. Oral Microbiol. 3, 5940. doi:10.3402/jom.v3i0.5940

22. Hyatt, D., Chen, G.-L., LoCascio, P.F., Land, M.L., Larimer, F.W., Hauser, L.J., 2010. Prodigal: prokaryotic gene recognition and translation initiation site identification. BMC Bioinform. 11, 119. doi:10.1186/1471-2105-11-119

23. Isaac, S., Flor-duro, A., Carruana, G., Puchades-carrasco, L., Quirant, A., Lopez-nogueroles, M., Pineda-lucena, A., Garcia-garcera, M., Ubeda, C., 2022. Microbiome-mediated fructose depletion restricts murine gut colonization by vancomycin-resistant Enterococcus. Nature Communications 7718. doi:10.1038/s41467-022-35380-5

24. Jangi, S., Gandhi, R., Cox, L.M., Li, N., Glehn, F. von, Yan, R., Patel, B., Mazzola, M.A., Liu, S., Glanz, B.L., Cook, S., Tankou, S., Stuart, F., Melo, K., Nejad, P., Smith, K., uolu, B. uuml m D.T. ccedil, Holden, J., kk, P.K. auml, Chitnis, T., Jager, P.L.D., Quintana, F.J., Gerber, G.K., Bry, L., Weiner, H.L., 2016. Alterations of the human gut microbiome in multiple sclerosis. Nature Communications 7, 1–11. doi:10.1038/ncomms12015

25. Kwilasz, A.J., Grace, P.M., Serbedzija, P., Maier, S.F., Watkins, L.R., 2015. The therapeutic potential of interleukin-10 in neuroimmune diseases. Neuropharmacology 96, 55–69. doi:10.1016/j.neuropharm.2014.10.020

26. Kursa MB, Rudnicki WR. Feature Selection with the Boruta Package. J. Stat. Softw. 2010;36(11).

27. Langmead, B., Salzberg, S.L., 2012. Fast gapped-read alignment with Bowtie 2. Nat. Methods 9, 357–359. doi:10.1038/nmeth.1923

28. Lassmann, H., 2017. Targets of therapy in progressive MS. Mult. Scler. J. 23, 1593–1599. doi:10.1177/1352458517729455

29. Leray, E., Yaouanq, J., Page, E.L., Coustans, M., Laplaud, D., Oger, J., Edan, G., 2010. Evidence for a two-stage disability progression in multiple sclerosis. Brain 133, 1900–1913. doi:10.1093/brain/awq076

30. Li, D., Luo, R., Liu, C.-M., Leung, C.-M., Ting, H.-F., Sadakane, K., Yamashita, H., Lam, T.-W., 2016. MEGAHIT v1.0: A fast and scalable metagenome assembler driven by advanced methodologies and community practices. Methods 102, 3–11. doi:10.1016/j.ymeth.2016.02.020

31. Li, M., Esch, B.C.A.M. van, Wagenaar, G.T.M., Garssen, J., Folkerts, G., Henricks, P.A.J., 2018. Pro- and anti-inflammatory effects of short chain fatty acids on immune and endothelial cells. European Journal of Pharmacology 831, 52–59. doi:10.1016/j.ejphar.2018.05.003

32. Lin, H., Peddada, S.D., 2020. Analysis of compositions of microbiomes with bias correction. Nat. Commun. 11, 3514. doi:10.1038/s41467-020-17041-7

33. Lublin, F.D., Reingold, S.C., Cohen, J.A., Cutter, G.R., Sørensen, P.S., Thompson, A.J., Wolinsky, J.S., Balcer, L.J., Banwell, B., Barkhof, F., Bebo, B., Calabresi, P.A., Clanet, M., Comi, G., Fox, R.J., Freedman, M.S., Goodman, A.D., Inglese, M., Kappos, L., Kieseier, B.C., Lincoln, J.A., Lubetzki, C., Miller, A.E., Montalban, X., O’Connor, P.W., Petkau, J., Pozzilli, C., Rudick, R.A., Sormani, M.P., Stüve, O., Waubant, E., Polman, C.H., 2014. Defining the clinical course of multiple sclerosis: The 2013 revisions. Neurology 83, 278–286. doi:10.1212/wnl.0000000000000560

34. Lukić, I., Getselter, D., Ziv, O., Oron, O., Reuveni, E., Koren, O., Elliott, E., 2019. Antidepressants affect gut microbiota and Ruminococcus flavefaciens is able to abolish their effects on depressive-like behavior. Transl. Psychiatry 9, 133. doi:10.1038/s41398-019-0466-x

35. Magoč, T., Salzberg, S.L., 2011. FLASH: fast length adjustment of short reads to improve genome assemblies. Bioinformatics 27, 2957–2963. doi:10.1093/bioinformatics/btr507

36. Meelu, P., Marin, R., Clish, C., Zella, G., Cox, S., Yajnik, V., Nguyen, D.D., Korzenik, J.R., 2014. Impaired Innate Immune Function Associated with Fecal Supernatant from Crohn’s Disease Patients. Inflammatory bowel diseases 20, 1139–1146. doi:10.1097/mib.0000000000000081

37. Mirza, A.I., Zhu, F., Knox, N., Forbes, J.D., Domselaar, G.V., Bernstein, C.N., Graham, M., Marrie, R.A., Hart, J., Yeh, E.A., Arnold, D.L., Bar-Or, A., O’Mahony, J., Zhao, Y., Hsiao, W., Banwell, B., Waubant, E., Tremlett, H., 2022. Metagenomic Analysis of the Pediatric-Onset Multiple Sclerosis Gut Microbiome. Neurology 98, e1050–e1063. doi:10.1212/wnl.0000000000013245

38. Miyake, S., Kim, S., Suda, W., Oshima, K., Nakamura, M., Matsuoka, T., Chihara, N., Tomita, A., Sato, W., Kim, S.-W., Morita, H., Hattori, M., Yamamura, T., 2015. Dysbiosis in the Gut Microbiota of Patients with Multiple Sclerosis, with a Striking Depletion of Species Belonging to Clostridia XIVa and IV Clusters. PLoS ONE 10, e0137429. doi:10.1371/journal.pone.0137429.s010

39. Nazareth, N., Magro, F., Appelberg, R., Silva, J., Gracio, D., Coelho, R., Cabral, J.M., Abreu, C., Macedo, G., Bull, T.J., Sarmento, A., 2015. Increased viability but decreased culturability of Mycobacterium avium subsp. paratuberculosis in macrophages from inflammatory bowel disease patients under Infliximab treatment. Medical Microbiology and Immunology 204, 647–656. doi:10.1007/s00430-015-0393-2

40. O’Connor, R.A., Anderton, S.M., 2008. Foxp3+ regulatory T cells in the control of experimental CNS autoimmune disease. J Neuroimmunol 193, 1–11. doi:10.1016/j.jneuroim.2007.11.016

41. Poles, M.Z., Juhász, L., Boros, M., 2019. Methane and Inflammation - A Review (Fight Fire with Fire). Intensiv. Care Med. Exp. 7, 68. doi:10.1186/s40635-019-0278-6

42. Pontieri, L., Greene, N., Wandall-Holm, M.F., Geertsen, S.S., Asgari, N., Jensen, H.B., Illes, Z., Schäfer, J., Jensen, R.M., Sejbæk, T., Weglewski, A., Mahler, M.R., Poulsen, M.B., Prakash, S., Stilund, M., Kant, M., Rasmussen, P.V., Svendsen, K.B., Sellebjerg, F., Magyari, M., 2024. Patterns and predictors of multiple sclerosis phenotype transition. Brain Commun. 6, fcae422. doi:10.1093/braincomms/fcae422

43. Quast, C., Pruesse, E., Yilmaz, P., Gerken, J., Schweer, T., Yarza, P., Peplies, J., Glöckner, F.O., 2013. The SILVA ribosomal RNA gene database project: Improved data processing and web-based tools. Nucleic Acids Research. doi:10.1093/nar/gks1219

44. Rahimlou, M., Hosseini, S.A., Majdinasab, N., Haghighizadeh, M.H., Husain, D., 2022. Effects of long-term administration of Multi-Strain Probiotic on circulating levels of BDNF, NGF, IL-6 and mental health in patients with multiple sclerosis: a randomized, double-blind, placebo-controlled trial. Nutr. Neurosci. 25, 411–422. doi:10.1080/1028415x.2020.1758887

45. Regen, T., Isaac, S., Amorim, A., Núñez, N.G., Hauptmann, J., Shanmugavadivu, A., Klein, M., Sankowski, R., Mufazalov, I.A., Yogev, N., Huppert, J., Wanke, F., Witting, M., Grill, A., Gálvez, E.J.C., Nikolaev, A., Blanfeld, M., Prinz, I., Schmitt-Kopplin, P., Strowig, T., Reinhardt, C., Prinz, M., Bopp, T., Becher, B., Ubeda, C., Waisman, A., 2021. IL-17 controls central nervous system autoimmunity through the intestinal microbiome. Sci. Immunol. 6. doi:10.1126/sciimmunol.aaz6563

46. Reynders, T., Devolder, L., Valles-Colomer, M., Remoortel, A.V., Joossens, M., Keyser, J.D., Nagels, G., D’hooghe, M., Raes, J., 2020. Gut microbiome variation is associated to Multiple Sclerosis phenotypic subtypes. Ann. Clin. Transl. Neurol. 7, 406–419. doi:10.1002/acn3.51004

47. Rukavishnikov, G., Leonova, L., Kasyanov, E., Leonov, V., Neznanov, N., Mazo, G., 2023. Antimicrobial activity of antidepressants on normal gut microbiota: Results of the in vitro study. Front. Behav. Neurosci. 17, 1132127. doi:10.3389/fnbeh.2023.1132127

48. Salami, M., Kouchaki, E., Asemi, Z., Tamtaji, O.R., 2019. How probiotic bacteria influence the motor and mental behaviors as well as immunological and oxidative biomarkers in multiple sclerosis? A double blind clinical trial. J. Funct. Foods 52, 8–13. doi:10.1016/j.jff.2018.10.023

49. Sawcer, S., Hellenthal, G., Pirinen, M., Spencer, C.C.A., Patsopoulos, N.A., Moutsianas, L., Dilthey, A., Su, Z., Freeman, C., Hunt, S.E., Edkins, S., Gray, E., Booth, D.R., Potter, S.C., Goris, A., Band, G., Oturai, A.B., Strange, A., Saarela, J., Bellenguez, C., Fontaine, B., Gillman, M., Hemmer, B., Gwilliam, R., Zipp, F., Jayakumar, A., Martin, R., Leslie, S., Hawkins, S., Giannoulatou, E., D’alfonso, S., Blackburn, H., Boneschi, F.M., Liddle, J., Harbo, H.F., Perez, M.L., Spurkland, A., Waller, M.J., Mycko, M.P., Ricketts, M., Comabella, M., Hammond, N., Kockum, I., McCann, O.T., Ban, M., Whittaker, P., Kemppinen, A., Weston, P., Hawkins, C., Widaa, S., Zajicek, J., Dronov, S., Robertson, N., Bumpstead, S.J., Barcellos, L.F., Ravindrarajah, R., Abraham, R., Alfredsson, L., Ardlie, K., Aubin, C., Baker, A., Baker, K., Baranzini, S.E., Bergamaschi, L., Bergamaschi, R., Bernstein, A., Berthele, A., Boggild, M., Bradfield, J.P., Brassat, D., Broadley, S.A., Buck, D., Butzkueven, H., Capra, R., Carroll, W.M., Cavalla, P., Celius, E.G., Cepok, S., Chiavacci, R., Clerget-Darpoux, F., Clysters, K., Comi, G., Cossburn, M., Cournu-Rebeix, I., Cox, M.B., Cozen, W., Cree, B.A.C., Cross, A.H., Cusi, D., Daly, M.J., Davis, E., Bakker, P.I.W. de, Debouverie, M., D’hooghe, M.B., Dixon, K., Dobosi, R., Dubois, B., Ellinghaus, D., Elovaara, I., Esposito, F., Fontenille, C., Foote, S., Franke, A., Galimberti, D., Ghezzi, A., Glessner, J., Gomez, R., Gout, O., Graham, C., Grant, S.F.A., Guerini, F.R., Hakonarson, H., Hall, P., Hamsten, A., Hartung, H.-P., Heard, R.N., Heath, S., Hobart, J., Hoshi, M., Infante-Duarte, C., Ingram, G., Ingram, W., Islam, T., Jagodic, M., Kabesch, M., Kermode, A.G., Kilpatrick, T.J., Kim, C., Klopp, N., Koivisto, K., Larsson, M., Lathrop, M., Lechner-Scott, J.S., Leone, M.A., Leppä, V., Liljedahl, U., Bomfim, I.L., Lincoln, R.R., Link, J., Liu, J., Lorentzen, A.R., Lupoli, S., Macciardi, F., Mack, T., Marriott, M., Martinelli, V., Mason, D., McCauley, J.L., Mentch, F., Mero, I.-L., Mihalova, T., Montalban, X., Mottershead, J., Myhr, K.-M., Naldi, P., Ollier, W., Page, A., Palotie, A., Pelletier, J., Piccio, L., Pickersgill, T., Piehl, F., Pobywajlo, S., Quach, H.L., Ramsay, P.P., Reunanen, M., Reynolds, R., Rioux, J.D., Rodegher, M., Roesner, S., Rubio, J.P., Rückert, I.-M., Salvetti, M., Salvi, E., Santaniello, A., Schaefer, C.A., Schreiber, S., Schulze, C., Scott, R.J., Sellebjerg, F., Selmaj, K.W., Sexton, D., Shen, L., Simms-Acuna, B., Skidmore, S., Sleiman, P.M.A., Smestad, C., Sørensen, P.S., Søndergaard, H.B., Stankovich, J., Strange, R.C., Sulonen, A.-M., Sundqvist, E., Syvänen, A.-C., Taddeo, F., Taylor, B., Blackwell, J.M., Tienari, P., Bramon, E., Tourbah, A., Brown, M.A., Tronczynska, E., Casas, J.P., Tubridy, N., Corvin, A., Vickery, J., Jankowski, J., Villoslada, P., Markus, H.S., Wang, K., Mathew, C.G., Wason, J., Palmer, C.N.A., Wichmann, H.-E., Plomin, R., Willoughby, E., Rautanen, A., Winkelmann, J., Wittig, M., Trembath, R.C., Yaouanq, J., Viswanathan, A.C., Zhang, H., Wood, N.W., Zuvich, R., Deloukas, P., Langford, C., Duncanson, A., Oksenberg, J.R., Pericak-Vance, M.A., Haines, J.L., Olsson, T., Hillert, J., Ivinson, A.J., Jager, P.L.D., Peltonen, L., Stewart, G.J., Hafler, D.A., Hauser, S.L., McVean, G., Donnelly, P., Compston, A., 2011. Genetic risk and a primary role for cell-mediated immune mechanisms in multiple sclerosis. Nature 476, 214–9. doi:10.1038/nature10251

50. Scazzone, C., Agnello, L., Sasso, B.L., Salemi, G., Gambino, C.M., Ragonese, P., Candore, G., Ciaccio, A.M., Giglio, R.V., Bivona, G., Vidali, M., Ciaccio, M., 2021. FOXP3 and GATA3 Polymorphisms, Vitamin D3 and Multiple Sclerosis. Brain Sci 11. doi:10.3390/brainsci11040415

51. Schmieder, R., Edwards, R., 2011. Quality control and preprocessing of metagenomic datasets. Bioinformatics 27, 863–864. doi:10.1093/bioinformatics/btr026

52. Shanmugavadivu, A., Carter, K., Zonouzi, A.P., Waisman, A., Regen, T., 2024. Protocol for the collection and analysis of the different immune cell subsets in the murine intestinal lamina propria. STAR Protoc. 5, 103154. doi:10.1016/j.xpro.2024.103154

53. Sumida, T.S., Cheru, N.T., Hafler, D.A., 2024. The regulation and differentiation of regulatory T cells and their dysfunction in autoimmune diseases. Nat. Rev. Immunol. 24, 503–517. doi:10.1038/s41577-024-00994-x

54. Talbot, J., Chow, H.H., Mahler, M., Buhelt, S., Hansen, R.H., Lundell, H., Vinther-Jensen, T., Hellem, M.N.N., Nielsen, J.E., Siebner, H.R., Essen, M.R. von, Sellebjerg, F., 2022. Relationship between cerebrospinal fluid biomarkers of inflammation and tissue damage in primary progressive multiple sclerosis. Mult Scler Relat Disord 68, 104209. doi:10.1016/j.msard.2022.104209

55. Tankou, S.K., Regev, K., Healy, B.C., Cox, L.M., Tjon, E., Kivisakk, P., Vanande, I.P., Cook, S., Gandhi, R., Glanz, B., Stankiewicz, J., Weiner, H.L., 2017. Investigation of probiotics in multiple sclerosis. Mult. Scler. J. 24, 58–63. doi:10.1177/1352458517737390

56. Tankou, S.K., Regev, K., Healy, B.C., Tjon, E., Laghi, L., Cox, L.M., Kivisäkk, P., Pierre, I.V., Hrishikesh, L., Gandhi, R., Cook, S., Glanz, B., Stankiewicz, J., Weiner, H.L., 2018. A probiotic modulates the microbiome and immunity in multiple sclerosis. Annals of Neurology 83, 1147–1161. doi:10.1002/ana.25244

57. Tremlett, H., Fadrosh, D.W., Faruqi, A.A., Hart, J., Roalstad, S., Graves, J., Spencer, C.M., Lynch, S.V., Zamvil, S.S., Waubant, E., Centers, U.N. of P.M., 2016. Associations between the gut microbiota and host immune markers in pediatric multiple sclerosis and controls. BMC neurology 16, 182–9. doi:10.1186/s12883-016-0703-3

58. Valenzuela, R.M., Costello, K., Chen, M., Said, A., Johnson, K.P., Dhib-Jalbut, S., 2007. Clinical response to glatiramer acetate correlates with modulation of IFN-gamma and IL-4 expression in multiple sclerosis. Mult Scler 13, 754–62. doi:10.1177/1352458506074510

59. Vinolo, M.A.R., Rodrigues, H.G., Hatanaka, E., Sato, F.T., Sampaio, S.C., Curi, R., 2011. Suppressive effect of short-chain fatty acids on production of proinflammatory mediators by neutrophils. The Journal of Nutritional Biochemistry 22, 849–855. doi:10.1016/j.jnutbio.2010.07.009

60. Waisman, A., Hauptmann, J., Regen, T., 2015. The role of IL-17 in CNS diseases. Acta Neuropathol. 129, 625–637. doi:10.1007/s00401-015-1402-7

61. Yogev, N., Bedke, T., Kobayashi, Y., Brockmann, L., Lukas, D., Regen, T., Croxford, A.L., Nikolav, A., Hövelmeyer, N., Stebut, E. von, Prinz, M., Ubeda, C., Maloy, K.J., Gagliani, N., Flavell, R.A., Waisman, A., Huber, S., 2022. CD4+ T-cell-derived IL-10 promotes CNS inflammation in mice by sustaining effector T cell survival. Cell Reports 38, 110565. doi:10.1016/j.celrep.2022.110565

62. Yoon, H., Gerdes, L.A., Beigel, F., Sun, Y., Kövilein, J., Wang, J., Kuhlmann, T., Flierl-Hecht, A., Haller, D., Hohlfeld, R., Baranzini, S.E., Wekerle, H., Peters, A., 2025. Multiple sclerosis and gut microbiota: Lachnospiraceae from the ileum of MS twins trigger MS-like disease in germfree transgenic mice—An unbiased functional study. Proc. Natl. Acad. Sci. 122, e2419689122. doi:10.1073/pnas.2419689122

63. Zanghì, A., Copetti, M., Avolio, C., Paolicelli, D., Romeo, M.A.L., Patti, F., Luca, G.D., Amato, M.P., Galgani, S., Sola, P., Salemi, G., Gallo, P., Granella, F., Romano, S., Zaffaroni, M., Bergamaschi, R., Pozzilli, C., Lus, G., Vianello, M., Trojano, M., D’Amico, E., Danni, M.C., Totaro, R., Lugaresi, A., Cocco, E., Valentino, P., Iaffaldano, P., Inglese, M., Bellantonio, P., Filippi, M., Rovaris, M., Clerici, V.T., Ferraro, D., Lus, G., Maniscalco, G.T., Morra, V.B., Pesci, I., Foschi, M., Aguglia, U., Montepietra, S., Marfia, G.A., 2025. Multiple sclerosis from onset to secondary progression: a 30-year Italian register study. J. Neurol., Neurosurg. Psychiatry 96, 1061–1069. doi:10.1136/jnnp-2025-335958

64. Zhou, X., Baumann, R., Gao, X., Mendoza, M., Singh, S., Sand, I.K., Xia, Z., Cox, L.M., Chitnis, T., Yoon, H., Moles, L., Caillier, S.J., Santaniello, A., Ackermann, G., Harroud, A., Lincoln, R., Gomez, R., Peña, A.G., Digga, E., Hakim, D.J., Vazquez-Baeza, Y., Soman, K., Warto, S., Humphrey, G., Farez, M., Gerdes, L.A., Oksenberg, J.R., Zamvil, S.S., Chandran, S., Connick, P., Otaegui, D., Castillo-Triviño, T., Hauser, S.L., Gelfand, J.M., Weiner, H.L., Hohlfeld, R., Wekerle, H., Graves, J., Bar-Or, A., Cree, B.A.C., Correale, J., Knight, R., Baranzini, S.E., 2022. Gut microbiome of multiple sclerosis patients and paired household healthy controls reveal associations with disease risk and course. Cell 185, 3467–3486.e16. doi:10.1016/j.cell.2022.08.021

