## Supplementary material for "Gut microbiome changes over the course of multiple sclerosis differentially influence autoimmune neuroinflammation": Suppl. Table 1

Supplementary Table 1: Characteristics of the participants in this study

| | | Non-MS | MS | S-MS | L-MS | *p* value |
|---|---|---|---|---|---|---|
| **Number of subjects** | | 26 | 25 | 11 | 14 | |
| **Gender - N(%)** | **male** | 12 (46,2%) | 6 (24,0%) | 3 (27,3%) | 3 (21,4%) | 0.144<br>(HLT vs MS patients)[1] |
| | **female** | 14 (53,8%) | 19 (76%) | 8 (72,7%) | 11 (78,6%) | |
| **Age (Min - Max)** | | 46 (19 - 69) | 44 (28 - 60) | 38 (28 - 51) | 49 (40 - 60) | 0.584<br>(HLT vs MS patients)[2] |
| **Disease duration in years (Min - Max)** | | | 17.0 (2 - 42) | 8.3 (2 - 15) | 23,8 (16 - 42) | |
| **EDSS - Median (inter quartile)** | | | 2.00 (1.00-3.25) | 1.00 (0-2.00) | 3.00 (1.50-3.62) | **0.0114***<br>(S-MS vs L-MS)[2] |
| **MSSS - Median (inter quartile)** | | | 1.69 (0.59-2.42) | 1.8 (0.53-2.01) | 1.67 (0.62-2.69) | 0.934<br>(S-MS vs L-MS)[2] |
| **DMT therapy (%)** | **Platform** | | 11 (44.0%) | 7 (63.7%) | 4 (28.6%) | |
| | **Oral** | | 3 (12.0%) | 2 (18.2%) | 1 (7.14%) | |
| | **Platform+Oral** | | 7 (28.0%) | 2 (18.2%) | 5 (35.7%) | |
| | **Platform+Oral+Monoclonal** | | 4 (16.0%) | 0 (0%) | 4 (28.6%) | |
| **Concomitant diseases** | **Depression** | 0 (0%) | 12 (48,0%) | 2 (18.2%) | 10 (71.4%) | **<0.0001******<br>(Non-MS vs MS patients)[1]<br>**0.0154***<br>(S-MS vs L-MS)[1] |
| | **Others*** | 13 (50%) | 4 (16.0%) | 2 (18.2%) | 2 (14.3%) | |

Non-MS, subjects without MS; S-MS, MS patients with short disease duration (1-15 years from onset); L-MS, MS patients with long disease duration (16 or more years from onset); DMD, disease modifying drugs (Platform: IFN-beta, Glatiramer acetate; Oral: Fumarate, Fingolimod; Monoclonal: Natalizumab); *dyslipidemia, hypertension, diabetes, thyroiditis, meningioma, rhinitis, hyperuricemia, asthma, cardiac disease; [1]Fisher's exact test; [2]Unpaired t test;
